# Functional optimization in distinct tissues and conditions constrains the rate of protein evolution

**DOI:** 10.1101/2024.07.22.604632

**Authors:** Dinara R. Usmanova, Germán Plata, Dennis Vitkup

**Author notes:** These authors contributed equally (Dinara R. Usmanova, Germán Plata). Correspondence to DV.

## Abstract

Understanding the main determinants of protein evolution is a fundamental challenge in biology. Despite many decades of active research, the molecular and cellular mechanisms underlying the substantial variability of evolutionary rates across cellular proteins are not currently well understood. It also remains unclear how protein molecular function is optimized in the context of multicellular species and why many proteins, such as enzymes, are only moderately efficient on average. Our analysis of genomics and functional datasets reveals in multiple organisms a strong inverse relationship between the optimality of protein molecular function and the rate of protein evolution. Furthermore, we find that highly expressed proteins tend to be substantially more functionally optimized. These results suggest that cellular expression costs lead to more pronounced functional optimization of abundant proteins, and that the purifying selection to maintain high levels of functional optimality significantly slows protein evolution. We observe that in multicellular species both the rate of protein evolution and the degree of protein functional efficiency are primarily affected by expression in several distinct cell types and tissues. Specifically, in developed neurons with upregulated synaptic processes in animals and in young and fast-growing tissues in plants. Overall, our analysis reveals how various constraints from the molecular, cellular, and species’ levels of biological organization jointly affect the rate of protein evolution and the level of protein functional adaptation.

## INTRODUCTION

Understanding protein molecular evolution and functional adaptation is a long-standing goal of biological research. The pioneering work of Zuckerkandl and Pauling (Zuckerkandl and Pauling 1962, 1965) demonstrated, to the initial surprise of many evolutionary biologists (Morgan 1998), that proteins accumulate amino acid substitutions in different lineages at approximately constant clock-like rate. The rate of protein evolution, i.e., the number of amino acid substitutions per protein site per unit time, varies by orders of magnitude across cellular proteins (Dickerson 1971; Koonin and Wolf 2010). But it is currently unclear what are the main biological mechanisms underlying this notable rate variability (Zhang and Yang 2015). Alongside elucidating the determinants of protein evolution, another key challenge in molecular and cell biology is understanding how and to what extent protein function is optimized in the context of different cell types and tissues. Previously, it has been observed that protein function, such as enzymatic efficiency, appears to be only moderately optimized in various species (Bar-Even, et al. 2011). But the origins of the diversity in functional optimization between proteins are not understood. Although the questions concerning the variability of protein evolutionary rates and protein functional optimization have been rarely considered together, they are likely to be closely intertwined. Because the majority of newly arising mutations are harmful to protein function (Futuyma and Kirkpatrick 2017), the strength of purifying selection against deleterious mutations, and therefore the rate of protein evolution, may depend on the level of optimized protein efficiency. Although plausible, the empirical evidence for this effect and its magnitude is currently lacking.

Multiple genomic and molecular correlates of protein evolutionary rates have been previously considered (Koonin and Wolf 2006; Rocha 2006), and the best known predictor is the level of protein expression (Pal, et al. 2001; Rocha and Danchin 2004; Pal, et al. 2006). The inverse relationship between protein expression and the rate of protein evolution, usually referred to as the Expression-evolutionary Rate (ER) correlation, shows that highly expressed proteins generally evolve slower than proteins with low expression levels; in other words, the sign of the ER correlation is negative. The level of gene expression explains up to a third of the protein evolutionary rate variance in various species, but the biological mechanisms underlying the ER correlation are not currently understood (Zhang and Yang 2015; Usmanova, et al. 2021). The ER correlation in animals is usually stronger in neural tissues (Drummond and Wilke 2008; Tuller, et al. 2008). However, the reasons for this interesting observation are not clear. At the molecular level, several models have been proposed to explain the ER correlation. One hypothesis proposed that ER is primarily mediated by increased protein stability necessary to prevent effects associated with toxic misfolding of highly expressed proteins (Drummond and Wilke 2008). However, multiple studies of various empirical datasets demonstrate only a small role played by differences in protein stability in explaining the variability of protein evolutionary rates (Plata and Vitkup 2018; Biesiadecka, et al. 2020; Usmanova, et al. 2021; Wu, et al. 2022).

The optimization of protein molecular function may not only slow the rate of protein evolution but may also underlie the ER correlation (Rocha 2006; Cherry 2010; Gout, et al. 2010). The total protein activity in the cell is usually proportional to the product of protein functional efficiency and expression level. Therefore, by optimizing protein efficiency organisms can express fewer copies of a protein while maintaining its total cellular activity. We refer to this mechanism as FORCE, as it is based on the idea of protein Functional Optimization to Reduce the Cost of Expression. According to FORCE, highly expressed proteins are under more stringent selection for functional optimization because improving their efficiency allows cells to save more resources required for protein production. Although computer simulations demonstrated how protein functional optimization, coupled with protein production costs, can in principle lead to the ER correlation (Cherry 2010), it is currently unclear whether this mechanism may explain a substantial fraction of the protein evolutionary rate variance and how the cost of expression in various tissues is related to protein functional optimization.

In this study we address several interrelated research questions raised above using analyses of multiple functional and genomics datasets. First, using data describing enzymes’ catalytic efficiency we investigate the role of functional optimization in constraining protein evolution. We then analyze comprehensive tissue- and cell-type-specific transcriptomics data to explore what biological and cellular processes are usually associated with strong ER correlations in animal and plant tissues. Next, we investigate the relationships between protein evolution, functional optimization, and expression patterns across tissues in multicellular organisms. Overall, our study reveals important biological mechanisms underlying the variability of protein evolutionary rates and demonstrates how specific molecular and cellular processes affect protein functional optimization.

## RESULTS

### Protein evolution and optimization of protein molecular function

To explore how the selection for functional optimality influences protein evolution it is necessary to consider a set of proteins with quantitative measurements of their functional efficiency. However, it is usually difficult to precisely characterize protein molecular function and to quantify its optimality across different proteins. Fortunately, based on the Enzyme Commission (EC) four-digit classification scheme (Webb 1992), diverse enzymatic functions have been well defined. Moreover, catalytic rates of many enzymes have been measured using accurate low throughput biochemical experiments (Chen and Vitkup 2007; Wittig, et al. 2018; Chang, et al. 2021). Two biochemical parameters, *k*_*cat*_ and *k*_*cat*_/*K*_*M*_, that characterize catalytic activities can be used to evaluate the functional efficiency of enzymes. The first order kinetic rate constant, *k*_*cat*_, quantifies the speed (turnover) of enzymatic reactions at saturating concentrations of substrates, and the specificity constant, *k*_*cat*_/*K*_*M*_, quantifies the second-order reaction rate at ligand concentrations substantially lower than the Michaelis constant, *K*_*M*_. Enzymes achieve their amazing catalytic efficiency primarily by stabilizing transition states of corresponding chemical reactions (Abeles, et al. 1992). However, depending on the chemical properties of substrates and the nature of catalyzed biochemical interconversions, it is much easier to achieve high kinetic rates for some enzymatic classes compared to others. This makes the comparison of absolute kinetic rates between different enzymatic classes not very informative. Therefore, following previous studies (Davidi, et al. 2018), we quantified the enzymatic functional optimality using relative catalytic rates. Specifically, we normalized absolute kinetic constants by the highest catalytic constants measured for enzymes from the same reaction class, i.e., enzymes sharing all four digits of the EC classification. The catalytic rates normalized in this way reflect the extent to which rate constants deviate from the maximal known rate from the same enzymatic class. To accurately estimate enzymatic efficiencies using the normalized rate constants we only considered EC classes with a certain minimal number of kinetic constant measurements available for different enzymes in each class (**Methods**). As we describe below, analyses based on the normalized kinetic rate constants 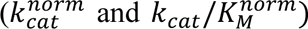 reveal insightful correlations between enzymatic optimality and protein evolutionary rates.

To analyze enzymatic functional optimality, we used a large collection of experimental *k*_*cat*_and *k*_*cat*_/*K*_*M*_measurements available in the Brenda (Chang, et al. 2021) and Sabio-RK databases (Wittig, et al. 2018). The largest number of kinetic constants were measured for *H. sapiens*, *A. thaliana*, and *E. coli* enzymes. Experimental measurements of kinetic constants from other species allowed us to estimate functional optimality for a substantial number of these enzymes. Notably, in all species we observed substantial correlations between the rate of protein evolution and functional optimality, quantified using either 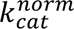 (Spearman’s *r* = −0.55, *p* = 1 · 10^-6^, for *H. sapiens*; *r* = −0.63, *p* = 8 · 10^-6^, for *A. thaliana*; *r* = −0.50, *p* = 1 · 10^-2^, for *E. coli*; **Fig. 1**) or 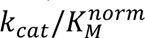 (**Supplementary fig. S1**). By analogy to the ER (expression-evolutionary rate) correlation, we refer to this correlation as KR, i.e., the correlation between the normalized kinetic rate (K) and the rate of protein evolution (R). The KR correlation, observed across a wide range (∼ 5 orders of magnitude) of protein optimality levels, demonstrates that higher optimality of protein molecular function indeed usually leads to substantially slower rates of protein evolution. Smaller evolutionary rates of proteins with highly optimized molecular function is likely a consequence of the additional constraints required to maintain optimal protein sequence, three-dimensional structure, and protein dynamics necessary for efficient function and catalysis (Konate, et al. 2019). We note that the explanatory power of functional optimality is substantially higher compared to multiple other protein biochemical properties, such as protein stability, solubility, and stickiness, which usually account for only a small percentage (1-5%) of the evolutionary rate variance (Plata, et al. 2010; Plata and Vitkup 2018; Usmanova, et al. 2021).

**Fig. 1.**
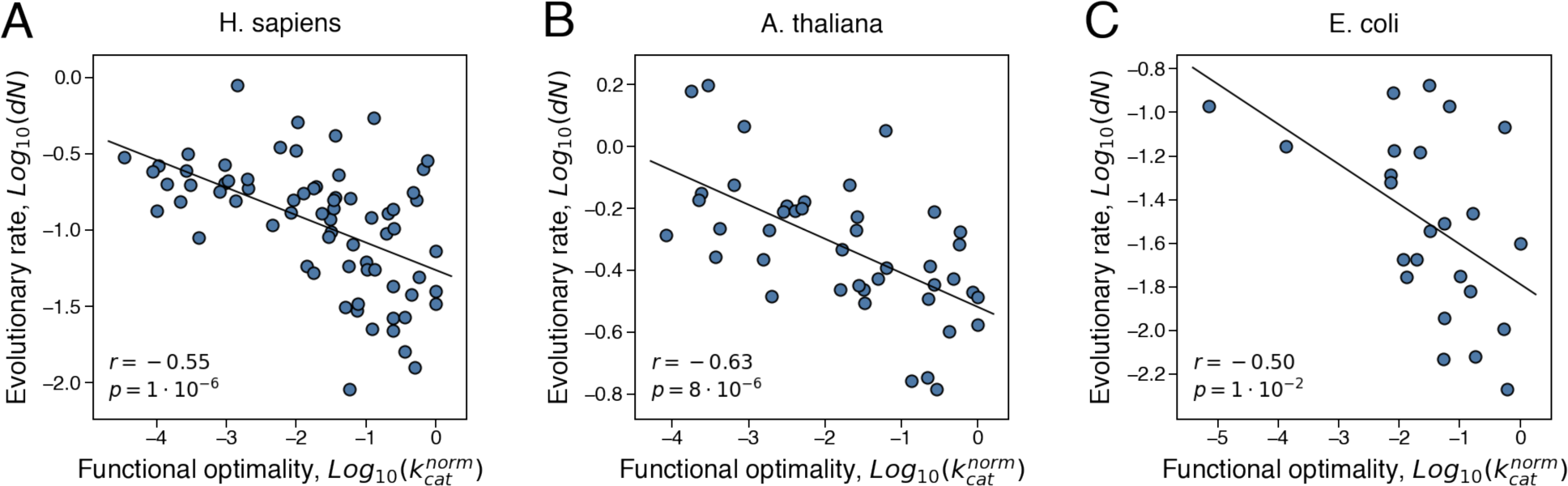
The correlation between protein functional optimality and the rate of protein evolution. Each point in the figures represents an enzyme from (**A**) *H. sapiens* (n=70), (**B**) *A. thaliana* (n=42), and (**C**) *E. coli* (n=24). Protein functional optimality was estimated using the normalized kinetic constant, 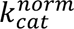, which quantifies the turnover catalytic rate relative to the maximal rate measured for the same reaction class. Evolutionary rate, *dN*, was calculated as the number of non-synonymous substitutions accumulated during the divergence of closely related orthologs per non-synonymous site. Spearman’s correlation coefficients and p-values are shown in each figure.

The evolutionary pressure to optimize protein function should be especially strong for highly expressed proteins, as such optimization allows cells to substantially reduce the number of expressed proteins. The FORCE model also suggests that in multicellular organisms the pressure for functional optimization and the ER correlation should both be stronger in the tissues with high protein production costs. Several possible scenarios can make certain tissues particularly sensitive to expression costs. One such scenario involves fast growing cells where protein expression is likely to be a major bottleneck. Constitutive protein expression, necessary due to constant protein turnover, is itself a major source of energy consumption (Buttgereit and Brand 1995; Rolfe and Brown 1997). Therefore, another potential scenario is either growing or non-growing cells with substantial and persistent energy needs. To explore these scenarios, we next investigated which tissues and cellular processes are typically associated with stronger ER correlations and with more pronounced protein functional optimization in multicellular organisms.

### The rate of protein evolution and gene expression in animals

Previous studies in animals demonstrated (Duret and Mouchiroud 2000; Zhang and Li 2004; Wang, et al. 2007) that proteins highly expressed in the brain generally evolve slowly (**Fig. 2A, Supplementary fig. S2A-C**). Thus, we first investigated to what extent expression in non-neural tissues further constrains protein evolutionary rates. We selected for this analysis several model organisms: *Homo sapiens* (Mele, et al. 2015), *Mus musculus* (Söllner, et al. 2017), *Drosophila melanogaster* (Leader, et al. 2018) and *Caenorhabditis elegans* (Spencer, et al. 2011); these organisms span ∼800 MYA divergence time, and their mRNA expression data are available across diverse tissues and cell types. Application of a multivariable regression analysis showed that only small additional fraction (<4%) of the protein evolutionary rate variance can be explained by considering expression in all tissues compared to expression in neural tissues only (**Supplementary Table 1**, **Methods**). Furthermore, we found that the strength of tissue-specific ER correlations in all species could be largely explained by the similarity between gene expression in a particular tissue and in the neural tissue with the strongest ER (**Fig. 2B, Supplementary fig. S2D-F**), confirming that the expression-based evolutionary constraints are primarily dominated by neural tissues. We next explored the expression breadth across tissues (Duret and Mouchiroud 2000; Park and Choi 2010), as this characteristic of global gene expression was suggested to be another important factor in constraining evolutionary rates in multicellular species (see **Methods**). We indeed found that the expression breadth correlates with evolutionary rates stronger than gene expression in many non-neural tissues, but substantially weaker than expression in neural tissues (**Supplementary Table 1**). The expression breadth explained little additional variance of evolutionary rates when combined with neural expression in the regression analyses (∼1% for all species). It has been previously demonstrated that protein expression is significantly correlated with the rate of protein polymorphisms in bacteria (Feugeas, et al. 2016). Similarly, based on the analysis of polymorphism data in the human population (Auton, et al. 2015), we observed that expression in neural tissues strongly correlates not only with the rates of interspecies protein evolution, but also with the per site frequency of polymorphisms across proteins (**Supplementary fig. S3**). This suggests that expression in neural tissues plays a dominant role in constraining protein evolution both between and within populations.

**Fig. 2.**
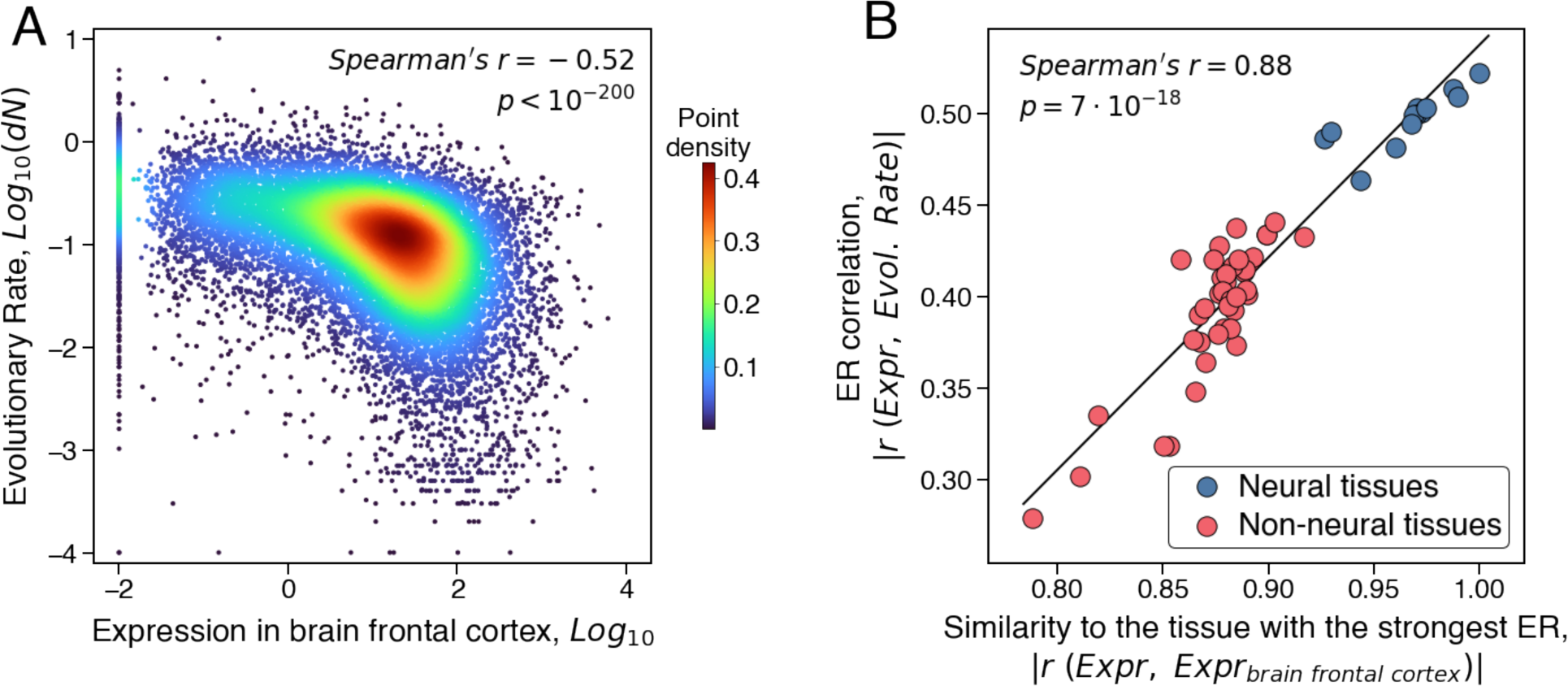
The relationship between expression in human tissues and the rate of protein evolution. (**A**) The correlation between gene expression in the brain frontal cortex and evolutionary rates of the corresponding human proteins. Evolutionary rates, *dN*, were calculated as the number of non-synonymous substitutions accumulated during the divergence of closely related orthologs per non-synonymous site (see Methods). Each point in the plot represents a human protein (n=18619), and the colors represent the point density. **(B)** The correlation between ER values across human tissues and the similarity of tissues’ genes expression to the frontal cortex; similar results were obtained for other animals (**Supplementary fig. S2D-F).** The similarity between tissues’ expression profiles was quantified using the Spearman’s correlation. Blue points (n=13) represent human neural tissues, and red points (n=40) represent non-neural tissues. The linear regression and the Spearman’s correlation coefficient were calculated based on all 53 tissues.

To further explore why the strongest ER correlation is observed in neural tissues, we first established that this effect is not primarily mediated by evolutionary properties of neural- or brain-specific genes. The removal of these genes from the analysis did not substantially weaken the ER correlation in neural tissues; for example, excluding 10% of the most neural-specific genes changed the explained evolutionary rate variance of the remaining genes by less than 1% in all species (**Methods**). Moreover, even for the subsets of genes most specific to non-neural tissues, the ER correlations based on their expression in the brain were almost always substantially stronger than based on expression in non-brain tissues (**Supplementary fig. S4, S5**; **Methods**). Strong ER correlations are also unlikely to result from the difference in the number of expressed genes in various tissues. Setting the abundance of lowly-expressed genes to zero to equalize the number of expressed genes across tissues decreased the explained evolutionary rate variance only by ∼0.5% in the neural tissues and also preserved the ranking of tissues by ER (Spearman’s *r* > 0.98, *p* < 4 · 10^-12^ for all species; **Methods**). Because it was previously demonstrated that the fraction of essential genes is similar across multiple mouse tissues (Cardoso-Moreira, et al. 2019), the observed differences in the ER correlation are also unlikely to originate from essentiality of brain-expressed genes. Therefore, the strong correlation between evolutionary rate and gene expression in neural tissues is likely to be mediated not by expression of brain-specific genes but by some inherent functional properties of neural cells that affect evolution of all proteins expressed in these cells.

To investigate the cellular properties that may underlie the ER correlations we took advantage of cell-type-specific brain expression data (Davie, et al. 2018; Saunders, et al. 2018; Zeisel, et al. 2018; Sugino, et al. 2019). We first used the single cell transcriptome dataset by Zeisel *et al*. (Zeisel, et al. 2018); the dataset covers hundreds of cell types including neurons and non-neurons across the entire mouse brain (**Methods**). Consistent with previous analyses (Hu, et al. 2020), we found that neurons generally have significantly stronger ER correlations than non-neuron brain cells (Mann-Whitney test *p* = 1 · 10^-20^; **Fig. 3A**); the ER correlations in central nervous system (CNS) neurons were significantly stronger than in neurons of the peripheral nervous system (PNS) (Mann-Whitney test *p* = 6 · 10^-10^). Notably, the strength of the ER correlation varies between different types of neurons, and we leveraged this variability to explore which specific cellular functions are usually upregulated in the neuron types associated with stronger ER correlations. To that end, we ranked all mouse genes based on how strongly their individual expression correlates with the ER strength across all CNS neuron types. We then used the gene set enrichment analysis (GSEA) (Subramanian, et al. 2005) to identify functional Gene Ontology (GO) categories that were associated with higher gene rankings (see **Methods**, **Supplementary Table 2**). The GSEA analysis showed that neurons with stronger ER correlations tend to have higher expression of genes associated with synaptic functions and related cellular processes (**Fig. 3B)**. An alternative GSEA approach, which ranked mouse genes based on the differential expression between CNS neuron types with high and with low ER strengths, implicated similar GO categories (**Methods**, **Supplementary Table 2**). Consistent with the GSEA results, we observed a strong correlation between the average expression of synaptic genes and the strength of ER correlation across neuron types (Spearman’s *r* = 0.87, *p* = 2 · 10^-42^; **Fig. 3C**). We again note that the correlation patterns identified by the GSEA analysis were not due to the synaptic genes themselves, as removing synapse-associated genes or genes from all upregulated GO categories from the ER calculations did not substantially change the GSEA results (**Methods**, **Supplementary Table 2**). Instead, it is likely that innate cellular properties of neurons with strongly upregulated synaptic functions make evolutionary rates of all proteins especially sensitive to expression levels in these cell types.

**Fig. 3.**
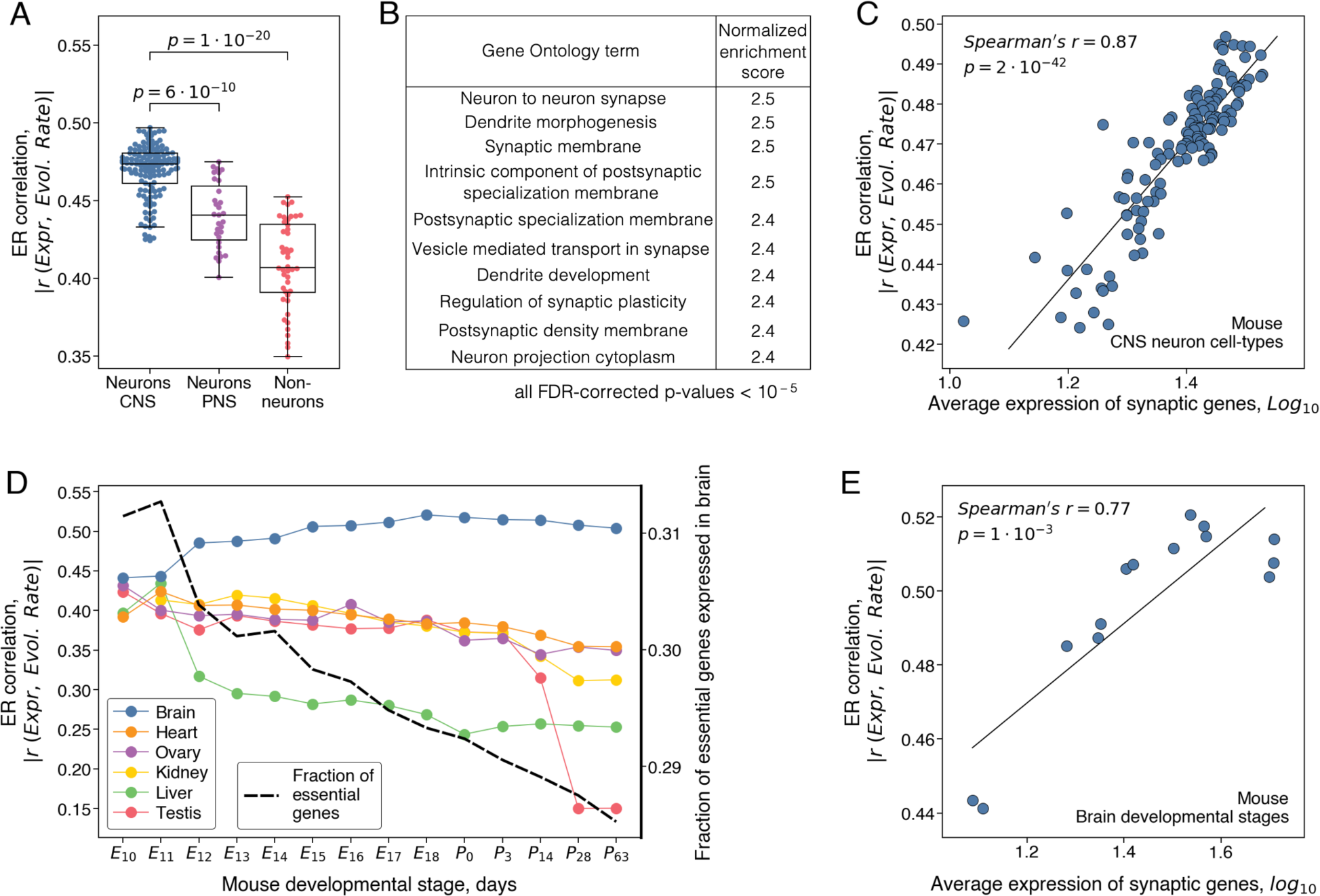
Functional properties of mouse brain cells and developmental stages that are associated with stronger ER correlations. (**A**) The strength of ER correlations across different cell types in the mouse nervous system (n=207). CNS neurons are shown in blue, PNS neurons in purple, and non-neuronal brain cells in red. The box plots show the median, the upper and lower quartiles of the ER strength, and the whiskers show the minimum and maximum values excluding outliers; P-values were calculated using the Mann-Whitney U-test. (**B**) The Gene Ontology (GO) terms associated with stronger ER in mouse CNS neurons. The 10 top GO terms ranked by the strength of the normalized enrichment score (see Methods) are shown; FDR-corrected P-values <10^-5^ for all presented GO terms. The complete list of significantly associated GO terms is provided in the **Supplementary Table 2**. (**C**) The correlation between the average expression of genes from the Synapse GO term (GO:0045202) and the cell-type specific ER strength. Each point represents a mouse CNS neuron type from (n=132). (**D**) The strength of ER correlation for mouse tissues across prenatal and postnatal developmental stages. Color lines represent different tissues, the X-axis shows developmental stages, and the left Y-axis shows the ER correlation strength. The right Y-axis shows the fraction of essential genes (represented by the dashed black line) expressed in the brain at different developmental stages. (**E**) The correlation between the average expression of genes from the Synapse GO term (GO:0045202) and the ER strength; the average gene expression and ER were calculated for the mouse brain at different developmental stages (n=14).

Next, we confirmed the correlation between the expression of synaptic genes and the ER strength using several independent datasets. First, we analyzed two additional comprehensive cell-type-specific transcriptomes of the mouse brain. One transcriptome was obtained using single-cell sequencing (Saunders, et al. 2018), while the other was obtained using the bulk RNA sequencing of cell populations distinguished by their genetic and anatomical markers (Sugino, et al. 2019). The GSEA analysis applied to these datasets confirmed the significant association between the upregulation of synapse-related functions and the strength of ER (**Supplementary fig. S6**, **Supplementary Table 2**). We also analyzed a region-specific gene expression dataset which covers the entire mouse brain (∼100 regions) and was obtained using *in situ* hybridization (Lein, et al. 2007). This dataset allowed us to relate the ER strength with brain regions’ physiological properties that were all measured in the same three-dimensional coordinate system (Erö, et al. 2018; Murakami, et al. 2018; Zhu, et al. 2018). Interestingly, we found that the ER strength was strongly correlated with the density of synapses (Zhu, et al. 2018) across the mouse brain (Spearman’s *r* = 0.66, *p* = 2 · 10^-13^), but not with the density of neurons (Erö, et al. 2018) (Spearman’s *r* = −0.051, *p* = 0.6) or the overall cellular density (Murakami, et al. 2018) (Spearman’s *r* = −0.15, *p* = 0.14) (**Supplementary fig. S7**). Finally, we analyzed single cell expression data from the *D. melanogaster* brain (Davie, et al. 2018), confirming that in the non-vertebrate species protein evolutionary rates also correlate significantly stronger with expression in neurons than in other brain cells (Mann Whitney test *p* = 9 · 10^-4^; **Supplementary fig. S6G**). The GSEA analysis applied to the *D. melanogaster* expression dataset also showed a significant association between the ER strength and the upregulation of synaptic and neuropeptide signaling functions, indicating the generality of these patterns in diverse species (**Supplementary fig. S6H**, **I**, **Supplementary Table 2**).

Functional properties of cells and their gene expression profiles vary not only between tissues and cell types but also across developmental stages, with especially rapid changes observed during embryogenesis. To investigate the changes in the strength of ER during development, we analyzed a temporal expression dataset that covers multiple mouse organs and includes both prenatal and postnatal developmental periods (Cardoso-Moreira, et al. 2019). In agreement with previous observations (Hu, et al. 2020), we found that expression in neural tissue becomes more strongly correlated with protein evolutionary rates with the progress of embryonic development (**Fig. 3D**). The maximal ER in the brain was reached around birth, without substantial further changes during the postnatal developmental stages. Qualitatively different behaviors were observed for non-neuronal tissues, in which the ER correlation monotonically decreased from the early to late developmental stages. The GSEA analysis performed across the brain development stages showed that the GO terms associated with stronger ER were generally similar to the terms associated with stronger ER in adult neurons (**Supplementary Table 2**). Specifically, we found that the ER strength strongly correlates with synaptic gene expression across the developmental stages (Spearman’s *r* = 0.75, *p* = 2 · 10^-3^; **Fig. 3E**). The patterns for neural and non-neural tissues were not due to the different number of genes expressed at various developmental stages, as ER correlations calculated using the same number of expressed genes showed a very similar temporal behavior with essentially the same ranking of samples by the ER strength (Spearman’s *r* = 0.99, *p* < 10^-20^, **Methods**). The strong ER correlation observed in the developed brain supports the conclusion that the ER variability between tissues is unlikely to be mediated by gene essentiality, as essential genes (Koscielny, et al. 2014) are more highly expressed in the brain during the early, rather than late, developmental stages (Cardoso-Moreira, et al. 2019) (**Fig. 3D**, dashed black line, **Methods**).

Overall, these results demonstrate that protein evolutionary rates in animals correlate more strongly with gene expression in developed neurons, especially in neurons with upregulated molecular and cellular functions related to synaptic activities. Our analysis also suggests that this effect is not primarily due to synaptic genes themselves, but that it is likely mediated by the functional properties of neurons in which synapse-related genes are highly expressed. Due to the generality of these results in animals, it is interesting to investigate cellular processes and functions that primarily affect protein evolutionary rates in multicellular organisms without neural tissues. Thus, we next considered the tissue-specific ER correlations and associated cellular processes in plants.

### The rate of protein evolution and gene expression in plants

We investigated the ER correlations in plants using multi-tissue RNA-seq data from three angiosperm species: *Zea mays* (corn) (Stelpflug, et al. 2016), *Arabidopsis thaliana* (Klepikova, et al. 2016), and *Glycine max* (soybean) (Shen, et al. 2014). As was reported previously, the similarity of plant tissues’ transcriptomes often reflects not only the relatedness of their morphological origins but also the similarity of their developmental stages (Klepikova, et al. 2016; Stelpflug, et al. 2016). We confirmed this observation in the considered plant species based on hierarchical clustering (**Methods**) of the tissues’ expression data (**Fig. 4**). This analysis resulted in distinct expression clusters representing samples from roots, leaves, stems, seeds, flowers, meristems, and also clusters that included multiple young or growing tissues of diverse origin. For example, the uppermost cluster in the dendrogram for corn (top in **Fig. 4A**) combines the samples from seedlings, root axes, leaf buds, developing seeds or flowers (see **Supplementary Table 3** for samples to cluster assignments). In contrast to animals, in plants we did not observe morphologically similar tissue types that have universally strong ER correlations (**Fig. 4**). However, plants’ samples with stronger ER tended to include young and growing tissues. Similar to animals, the observed patterns were not primarily due to the genes specific to growing plant tissues, as removal of such genes did not substantially affect the strength of ER (**Methods**); for example, removing 10% of the genes most specific to the growing tissues changed the fraction of the evolutionary rate variance explained by expression in the plant tissues with strong ER by less than 1%. This again suggests that it is the functional properties of fast-growing plant tissues that likely make evolutionary rates of plant proteins especially sensitive to expression in the corresponding cells and organs.

**Fig. 4.**
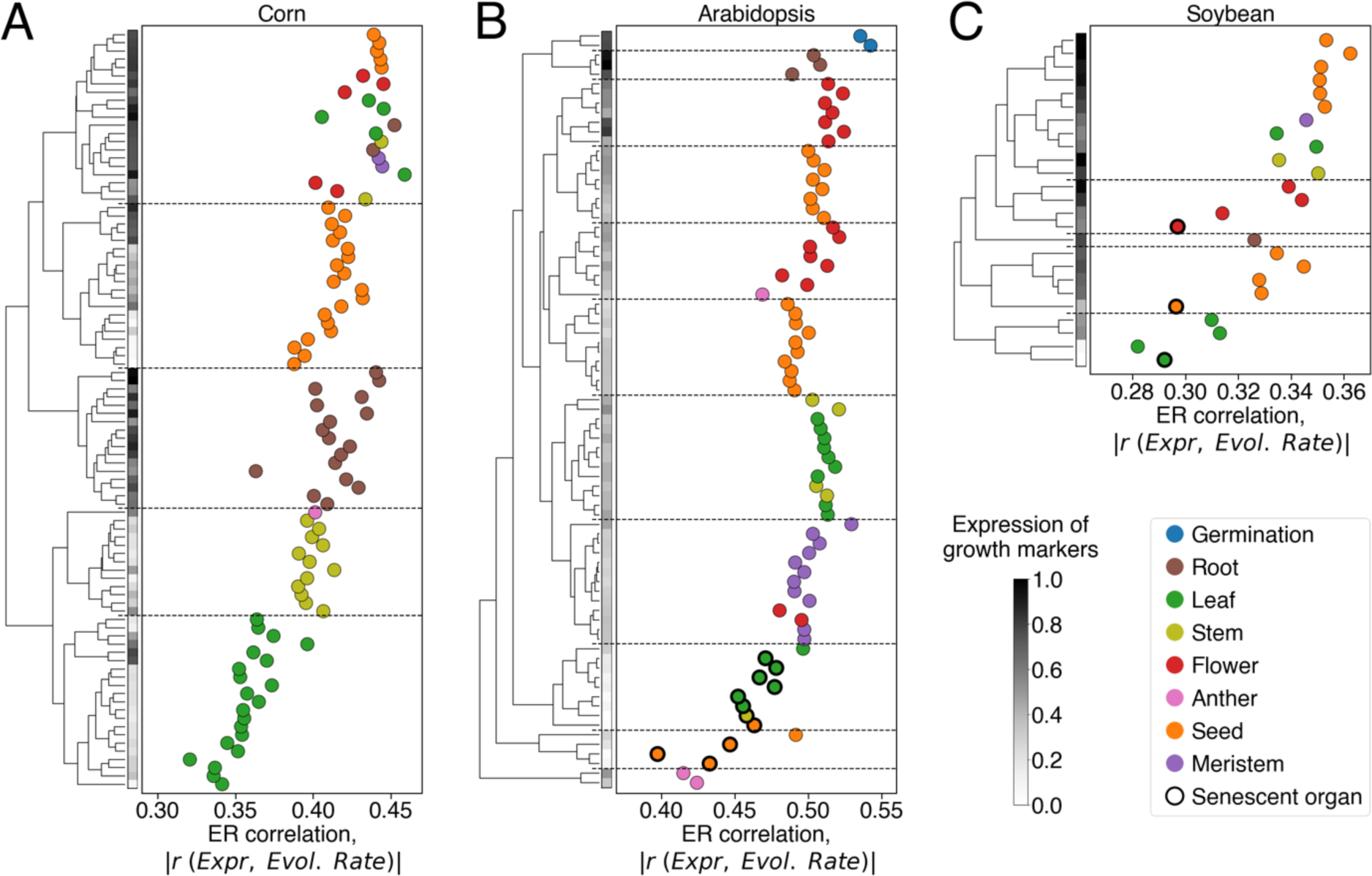
The relationship between expression in plant tissues and the rate of protein evolution. The data are shown for transcriptomes of (**A**) corn (Stelpflug, et al. 2016), (**B**) Arabidopsis (Klepikova, et al. 2016), and (**C**) soybean (Shen, et al. 2014). Each point in the figure represents a plant tissue sample, point colors represent different plant tissue types described in the legend; senescent samples are shown as black-edged circles. The X-axis represents the strength of the ER correlation. The left panel of each plot shows the hierarchical clustering dendrogram of the plant transcriptomes, with the clustering distance metric calculated as one minus the squared Pearson’s correlation between samples’ expression profiles. The horizontal dashed lines separate major clusters of the dendrogram. The vertical grey scale colormaps on the left side of the plots show the scaled average expression of cell growth markers in the corresponding plant tissues. The genes strongly upregulated in fast growing root cells (Huang and Schiefelbein 2015) were used as the growth markers for Arabidopsis (see **Methods**); the orthologs of the Arabidopsis growth markers were used as the growth markers for corn and soybean.

To understand the functional properties of plant cells associated with strong ER, we again used the GSEA enrichment analysis. In all three considered plant species we found that similar GO terms are usually associated with stronger ER correlations (**Supplementary fig. S8**, **Supplementary Table 2**). These upregulated GO terms primarily represent growth-related functional categories, such as translation, cell wall biosynthesis, and microtubule cytoskeleton organization/movement (**Fig. 5A**). The implicated functional categories suggest that strong ER correlations in plants are associated with cellular elongation and growth. The growth of plant organs is known to be initiated in the zone of undifferentiated meristematic cells and consists of three consecutive stages: cells division, elongation and differentiation (Taiz, et al. 2015). Therefore, we used the gene markers of these growth stages (Huang and Schiefelbein 2015) to further investigate how the marker’s average expression correlates with ER across tissues (**Methods**). This analysis indeed demonstrated that expression of the cellular elongation markers most strongly correlates with the strength of ER in all three plant species (Spearman’s *r* = 0.72, *p* = 8 · 10^-16^, for corn; *r* = 0.75, *p* = 3 · 10^-15^, for Arabidopsis; *r* = 0.83, *p* = 4 · 10^-7^, for soybean; **Fig. 5B, C, D**). The expression of the cell division markers showed a weaker and less significant correlation with the ER strength (Spearman’s *r* = 0.48, *p* = 1 · 10^-6^, for corn; *r* = 0.15, *p* = 0.2, for Arabidopsis; *r* = 0.36, *p* = 0.08, for soybean; **Supplementary fig. S9**). Finally, the association between the cell differentiation markers and ER was either not significant or was significant in the direction opposite to the other two markers (Spearman’s *r* = −0.63, *p* = 2 · 10^-11^, for corn; *r* = −0.02, *p* = 0.9, for Arabidopsis; *r* = −0.32, *p* = 0.1 for soybean), confirming a substantial decrease of the ER strength for plant tissues entering the differentiation stage.

**Fig. 5.**
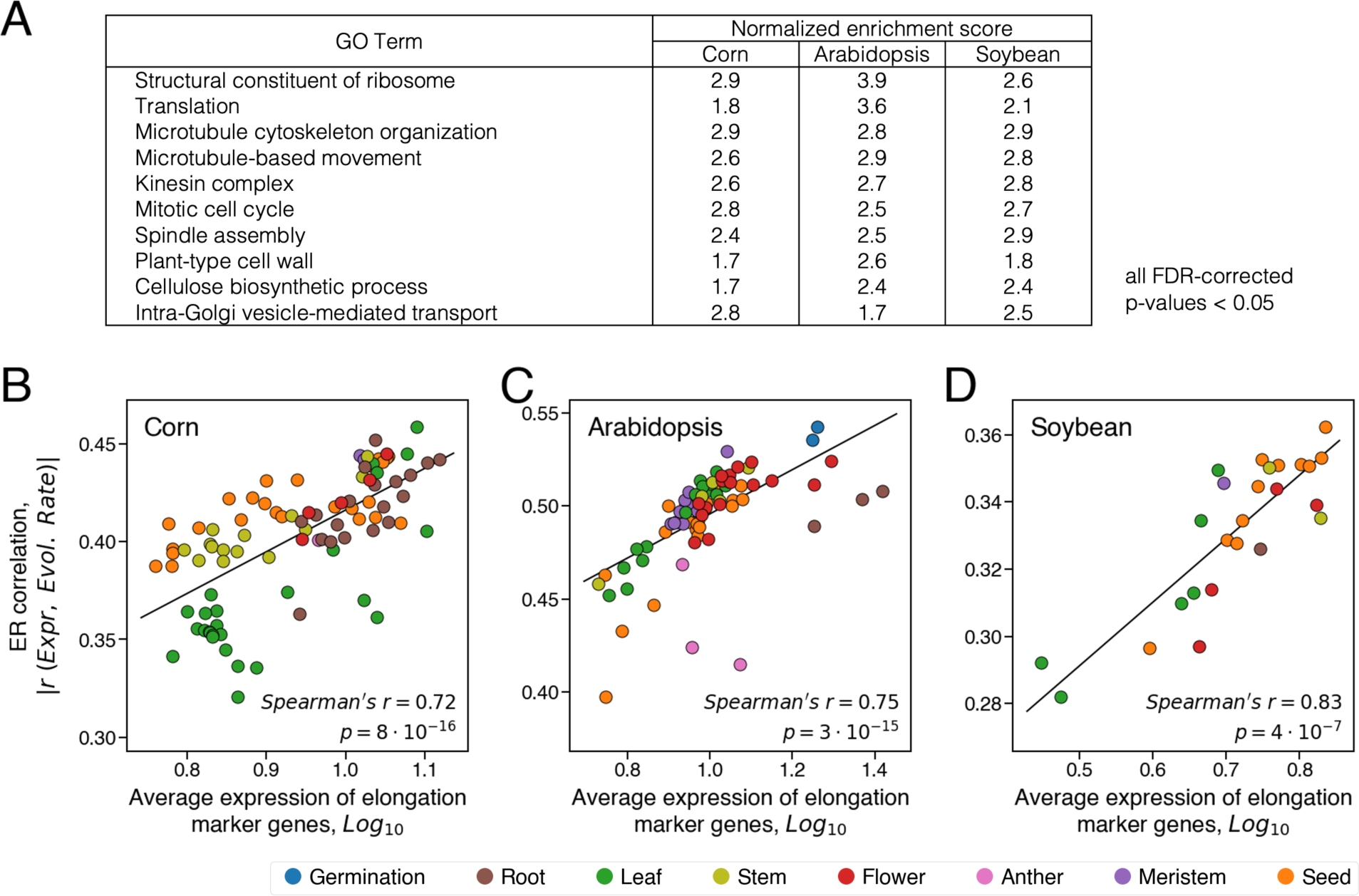
Functional properties of plant tissues that are associated with stronger ER correlations. **(A)** GO terms significantly associated with tissues showing stronger ER correlations in the three considered plant species, corn, Arabidopsis, and soybean. Normalized enrichment scores for 10 representative GO terms are shown; FDR-corrected P-values <0.05 for all presented GO terms. The complete list of upregulated GO terms is provided in the **Supplementary Table 2**. (**B**, **C**, **D)** The relationship between the average expression of elongation growth markers and the tissue-specific ER strength for (**B**) corn (n=92), (**C**) Arabidopsis (n=79), and (**D**) soybean (n=25). Each point in the figure represents a plant tissue, and point colors represent the plants’ tissue types described in the legend.

While different cellular functions are associated with strong ER correlations in animals and plants, our results suggest that the strongest correlations are usually observed in tissues with high expression costs. In plants, our analysis implicates cells from various tissues that are rapidly growing, and therefore likely prioritizing their carbon and energy resources for novel protein production. In animals, synaptic activities in the brain require a substantial and persistent energy supply (Harris, et al. 2012). Thus, it may be particularly important to reduce expression burden for neurons with high densities of synaptic connections. Having identified the tissues with the strongest ER correlations, we next investigated to what extent the correlation between protein functional optimality and evolutionary rate may mediate the ER correlation and its variability across tissues.

### The role of protein functional optimization in mediating the ER correlation

To investigate the relationship between the optimization of protein function and the ER correlation, we considered the sets of *H. sapiens*, *A. thaliana,* and *E. coli* enzymes with estimated levels of their functional optimality (**Fig. 1**). First, we confirmed that for the proteins from these sets the ER correlation is significant and similar in strength to the ER correlation for the whole corresponding proteomes (Spearman’s *r* = −0.64, *p* = 3 · 10^-9^, for *H. sapiens*; *r* = −0.61, *p* = 2 · 10^-5^, for *A. thaliana*; *r* = −0.75, *p* = 3 · 10^-5^, for *E. coli*, **Supplementary fig. S10**). This result suggests that the mechanisms underlying the whole-proteome ER correlation play a similar role in the evolution of these proteins. The FORCE mechanism proposes that highly expressed proteins are more functionally optimized to relieve the burden associated with their production costs. Consistent with this model, we observed in all three species significant correlations between protein expression and functional optimality (Spearman’s *r* = 0.52, *p* = 4 · 10^-6^, for *H. sapiens*; *r* = 0.45, *p* = 3 · 10^-3^, for *A. thaliana*; *r* = 0.46, *p* = 2 · 10^-2^, for *E. coli*; **Fig. 6A, B, C**). By analogy to the ER and KR correlations, we refer to this revealing correlation as EK, i.e., the correlation between expression (E) and kinetic constants (K).

**Fig. 6.**
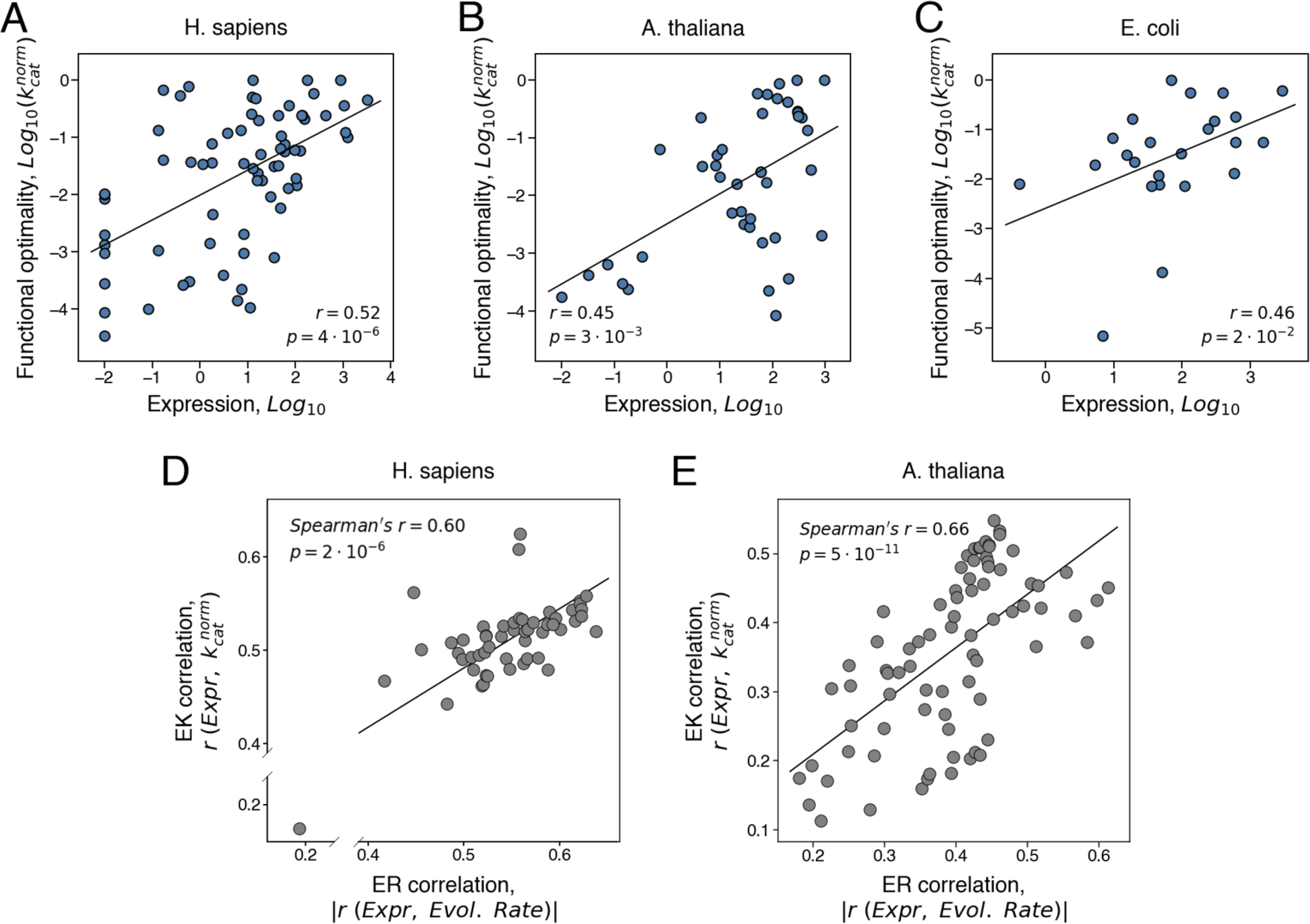
The relationship between the level of protein functional optimality, expression, and the rate of protein evolution. (**A, B, C**) The correlation between protein expression level and protein functional optimality (the EK correlation). Each point on the plots represents an enzyme from (**A**) *H. sapiens* (n=70), (**B**) *A. thaliana* (n=42), and (**C**) *E. coli* (n=24). Protein functional optimality was estimated using the normalized kinetic constant, 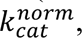 which quantifies the turnover catalytic rate relative to the maximal rate measured for the same reaction class. For multicellular species, the expression in the tissue with the strongest ER for enzymes was used (the brain basal ganglia for *H. sapiens* and the seedling root for *A. thaliana*). (**D**, **E**) The relationship between ER correlation (the correlation between expression and evolutionary rate) and the EK correlation (the correlation between expression and protein functional optimality, quantified as 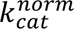. Each point represents a tissue from (**D**) *H. sapiens* (53 tissues) or (**E**) *A. thaliana* (79 tissues); the strength of ER and EK correlations were quantified in different tissues using the Spearman’s correlation.

Our analyses of protein expression across tissues demonstrated that the rate of protein evolution is especially sensitive to expression in several specific cell types and tissues, such as neurons in animals (**Fig. 2**) and actively growing tissues in plants (**Fig. 4**). The tissues most sensitive to expression costs are likely to exert the highest selective pressure for the protein functional optimization. As a result, both the EK and ER correlations should be stronger in tissues with high expression costs and weaker in other tissues. In agreement with this prediction, in both animals and plants we observed significant correlations between the strengths of ER and EK across tissues (Spearman’s *r* = 0.60, *p* = 2 · 10^-6^, for *H. sapiens*; *r* = 0.66, *p* = 5 · 10^-11^, for *A. thaliana*; **Fig. 6D, E**), with the highest correlations observed in the brain for *H. sapiens* and fast-growing tissues for *A. thaliana*.

Finally, we quantitatively investigated what fraction of the ER correlation can be explained by the variability in functional optimality. To that end, we used the semi-partial correlations to calculate the unique and shared contributions of independent variables, in this case expression and protein functional optimality, to explaining the variance of protein evolutionary rates. This analysis demonstrated that after controlling for protein functional optimality, quantified using 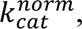 the evolutionary rate variance explained by expression substantially decreases: by ∼1/2 for *H. sapiens* (from 41% to 17%), ∼2/3 for *A. thaliana* (from 38% to 14%), and ∼1/3 for *E. coli* (from 56% to 34%); similar results were obtained for the multicellular species when controlling for 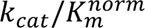 (**Supplementary fig. S11**). The observed decreases in the variance explained correspond to the shared fractional contribution of protein expression and functional optimality to the variance of evolutionary rate, and the remaining fractions represent their unique contributions (**Fig. 7**). The large explanatory effect sizes of KR are especially remarkable because our 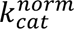-based estimations of protein optimality rely on a simple and heuristic normalization procedure. We note that the fractions of ER unexplained by 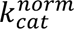 do not necessarily indicate that they are unrelated to protein function. Kinetic constants, such as *k*_*cat*_, are clearly not the only parameters optimized in protein evolution. The refinements of various other functional properties, such as the efficiency of allosteric regulation, covalent modification, and protein-protein binding, may further increase the fraction of ER explained by functional optimality. Taken together, our results demonstrate not only an important role played by the optimization of protein efficiency in constraining protein evolution, but also its major role in mediating the ER correlation.

**Fig. 7.**
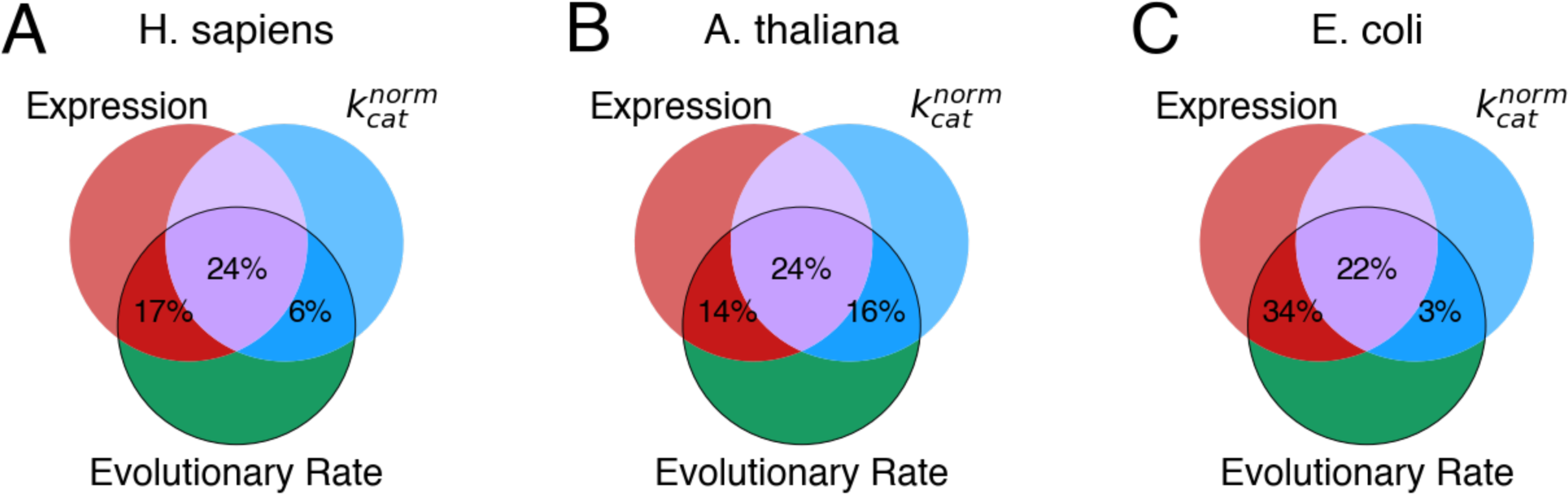
The fractions of the evolutionary rate variance explained by protein functional optimality and expression. The Venn diagram shows for (**A**) *H. sapiens*, (**B**) *A. thaliana*, and (**C**) *E. coli*, the fractions of the evolutionary rate variance explained by expression and protein functional optimality; functional optimality was quantified using the normalized kinetic constant 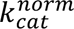. The two-way intersections (red and blue) represent the unique contributions of expression and functional optimality, respectively, and three-way intersection (purple) represents the shared contribution of these two factors. For multicellular species, the expression in the tissue with the strongest ER for enzymes was used (the brain basal ganglia for *H. sapiens* and the seedling root for *A. thaliana*). The unique and shared contributions were estimated using semi-partial correlations (see **Methods**).

## DISCUSSION

The presented results demonstrate that maintaining optimal protein function, for example high values of enzyme kinetic constants, imposes substantial constraints on protein sequences. This effect decreases the rate of amino acid substitutions and thus slows protein evolution. Our analysis of empirical data shows that the variability in functional optimality explains a substantial fraction (30-40%) of the protein evolutionary rate variance in such diverse organisms as *H. sapiens*, *A. thaliana*, and *E. coli* (the KR correlation, **Fig. 1**). We note that in addition to the optimization of protein function other protein- and structure-specific factors are likely to influence the rate of protein evolution (Wolf, et al. 2008; Wolf, et al. 2010).

Our results also suggest that the functional model of protein evolution (**Fig. 8**), based on the KR correlation and the FORCE mechanisms, may explain up to half of the correlation between the rate of protein evolution and protein expression. In addition to the KR correlation, we found that protein expression and functional efficiency also correlate between themselves across tissues (the EK correlation, **Fig. 6A-C**). The EK correlation likely emerges because both protein efficiency and expression level tend to increase together to meet the cellular demand for the total protein activity. Because protein expression is usually associated with certain cellular costs (Dekel and Alon 2005; Plata, et al. 2010; Kafri, et al. 2016), increasing the total protein activity exclusively through upregulation of protein expression is disadvantageous. On the other hand, the functional optimization of protein sequence is limited by the entropic factor, i.e., there are many more sequences with sub-optimal than with optimal function. Balancing between the expression cost and the mutation-selection balance for the functional optimization, the cell then satisfies the activity demand via both avenues simultaneously. As a result, proteins with high total activity demand tend to display both high expression and high functional efficiency, while proteins with low demand tend to have low expression and low efficiency. Because protein expression level positively correlates with functional optimality, and functional optimality reduces the rate of protein evolution, highly expressed proteins usually evolve slowly, i.e., display the ER correlation (**Fig. 8**). Our results show that protein functional optimality and evolutionary rate are primarily affected by expression in the same tissues, likely to be the ones most sensitive to expression costs (**Fig. 6D, E**).

**Fig. 8.**
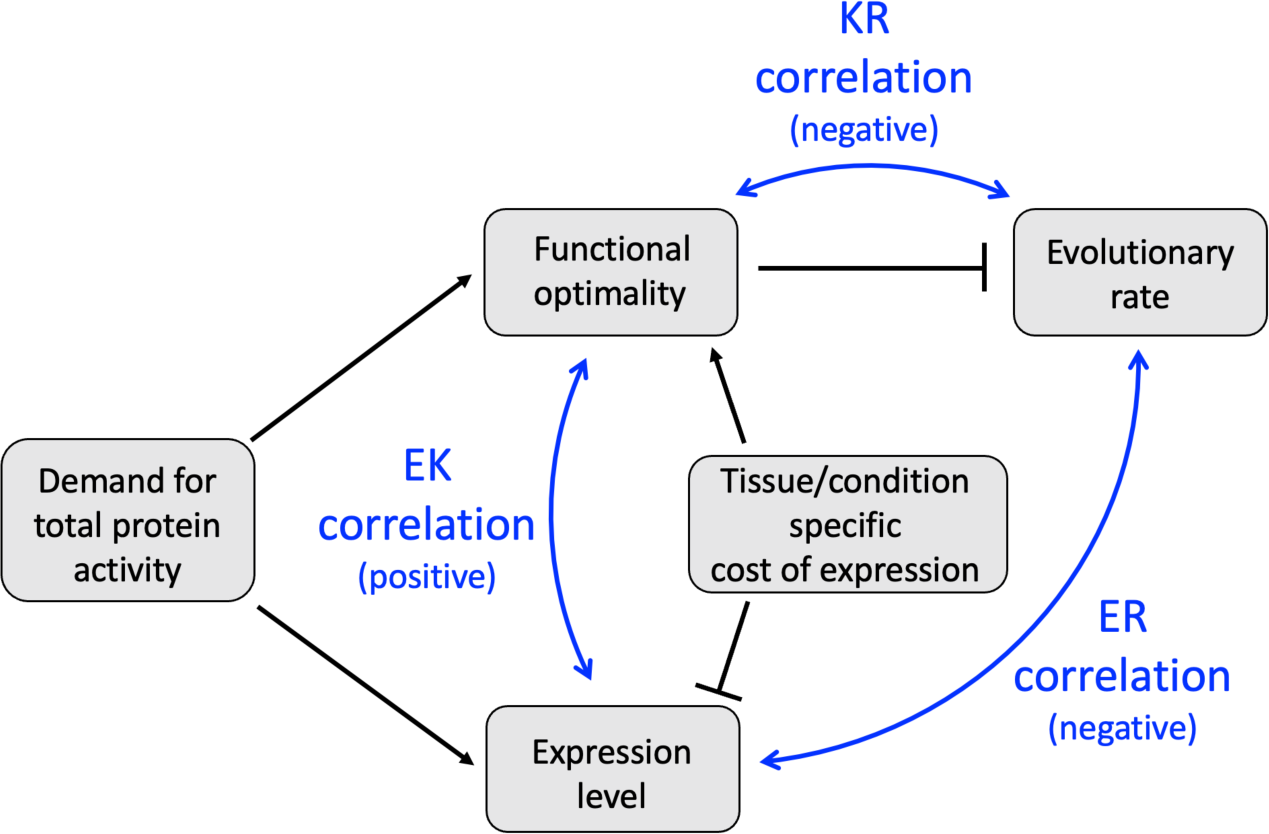
Functional model of protein evolution. The figure illustrates the functional model of protein evolution and the Functional-Optimization-to-Reduce-the-Cost-of-Expression (FORCE) mechanism that are supported by the presented data. Grey rectangles represent molecular and cellular properties and processes. Black lines indicate stimulatory (arrows) and inhibitory (T-bars) effects. The bidirectional blue arrows indicate the key experimentally observed correlations: the negative ER correlation between expression and evolutionary rate, the negative KR correlation between protein functional optimality and evolutionary rate, and the positive EK correlation between expression and protein functional optimality. A higher demand for the total protein activity simultaneously leads to higher protein expression and increased functional optimality (the EK correlation). The requirement to maintain high functional optimality, in turn, constrains protein sequence and slows the rate of protein evolution (the ER correlation). Furthermore, particularly high costs of protein expression in certain tissues and conditions facilitates the optimization of protein functions and leads to stronger EK and ER correlations in the tissues most sensitive to expression costs. We note that the FORCE mechanism does not generally depend on the nature of the protein expression cost, and different expression costs, or their various combinations, can lead to similar patterns of the ER and EK correlations.

We find that in multicellular organisms the strength of the ER correlation and the pressure to optimize protein function are usually associated with upregulation of distinct cellular processes, such as synaptic activities in animals (**Fig. 3**) and growth-related processes in plants (**Fig. 5**). Functional optimization may help to relieve the expression burden in plant tissues with rapid cellular growth and active translation. Fast-growing plant cells experience a high demand for ATP and carbon required for biosynthesis. For example, it was estimated that in *A. thaliana* fast-growing leaves spend ∼40% of their ATP on protein production, while slow-growing leaves spend about 3 times less (Li, et al. 2017). Similarly, energetic and morphological properties of animal neurons, especially neurons with upregulated synaptic activities, are likely to result in particularly high costs of protein expression. Multiple evidence suggest that the brain and neurons are highly sensitive to energy limitations, with the majority (up to ∼80%) of neuronal ATP spent on synaptic repolarization (Alle, et al. 2009; Harris, et al. 2012; Magistretti and Allaman 2015). In addition to substantial energy consumption, large volumes of neuronal dendritic trees may provide a protein trafficking burden as protein expression primarily takes place in the soma (Maday, et al. 2014). While one mechanism to reduce expression costs in neurons is to optimize protein function, another general mechanism is to decrease the rate of protein turnover. The rate of protein turnover is indeed substantially slower in the brain compared to other tissues (Fornasiero, et al. 2018), and it is slower in neurons compared to glial cells (Dörrbaum, et al. 2018). Slower evolution of proteins highly expressed in the brain may also lead to slower evolution of related cellular systems and properties. For example, it has been demonstrated that cellular transcriptome (Brawand, et al. 2011; Chen, et al. 2019), metabolome (Ma, et al. 2015), and tissue-specific codon usage (Plotkin, et al. 2004) evolve significantly slower in the brain compared to other tissues.

Proteins, cells, and tissues of multicellular organisms do not function in isolation, but rather as an integrated system that insures proper physiological responses and species’ survival. Therefore, it is of keen interest to understand the optimization of individual biological components, such as proteins, in the context of complex biological systems. We and others have previously investigated the influence of cellular protein-protein and metabolic networks on the evolution of individual proteins (Fraser, et al. 2002; Vitkup, et al. 2006). Our present work reveals the striking variability of protein functional optimization and how that variability affects protein evolution. We hope that our study will be an important step towards understanding the interdependence between protein functional optimization, protein evolution, and expression across tissues in multicellular organisms.

## METHODS

### Gene expression datasets used in the analyses

The following transcriptomes were used in this study: tissue-specific transcriptomes of *Homo sapiens* (Mele, et al. 2015), *Mus musculus* (Söllner, et al. 2017), *Drosophila melanogaster* (Leader, et al. 2018), *Caenorhabditis elegans* (Spencer, et al. 2011)*, Arabidopsis thaliana* (Klepikova, et al. 2016), *Zea mays* (Stelpflug, et al. 2016) and *Glycine max* (Shen, et al. 2014); cell type-specific transcriptomes of the brain of *M. musculus* (Saunders, et al. 2018; Zeisel, et al. 2018; Sugino, et al. 2019) and *D. melanogaster* (Davie, et al. 2018); region-specific transcriptome of *M. musculus* brain (Lein, et al. 2007); tissue-specific transcriptome at different developmental stages of *M. musculus* (Cardoso-Moreira, et al. 2019), and the transcriptome of *Escherichia coli* measured at log phase growth (McClure, et al. 2013). Data sources, number of samples, sequencing technique, and specific details on data extraction for each dataset are provided in **Supplementary Table 4**. In our analyses, we only used expression levels for the chromosomal protein-coding genes. For all single cell datasets, we also excluded cell-type clusters with low expression resolution and used only clusters with at least 200,000 UMI counts. We TPM-normalized tissue-specific transcriptomes for *H. sapiens*, *M. musculus*, *D. melanogaster*, *C. elegans*, and *E. coli* before subsequent analyses. We used the trimmed mean of M-values (TMM) normalization method (Robinson and Oshlack 2010) available in the edgeR package (Robinson, et al. 2010) for all plant tissue-specific transcriptomes and for all animal single-cell transcriptomes to allow comparisons across samples in analyses such as the GSEA. All expression datasets used in this work are available in the **Supplementary Table 5**.

### Calculation of evolutionary rates

We used the rate of non-synonymous substitutions, *dN*, as a measure of protein evolutionary rate. To calculate *dN* values for pairs of orthologous proteins, we utilized the PAML package (Yang 1997). We identified orthologous proteins as bidirectional best hits in pairwise local alignments between proteins from two species. The pairwise alignments were generated using Usearch (Edgar 2010), and included only protein pairs for which the corresponding alignments had E-value < 10^-6^, were at least 30 amino acids long, and covered at least 70% of the length of both proteins. The orthologous pairs of species used in our analysis were: *Mus musculus* – *Homo sapiens*, *Drosophila melanogaster* – *Drosophila yakuba*, *Caenorhabditis elegans* – *Caenorhabditis briggsae*, *Zea mays* (corn) – *Oryza sativa* (rice), *Glycine max* (soybean) – *Medicago truncatula*, and *Escherichia coli* - *Salmonella enterica*. The CDS sequences for the proteins in each species were obtained from the Ensemble database (Cunningham, et al. 2022).

For *H. sapiens* and *A. thaliana* we used a multi-species approach to obtain accurate estimates of protein evolutionary rate. Notably, the standard procedure which uses only a pair of closely related species, did not find reliable orthologs for a fraction of proteins, thus precluding us from inferring their evolutionary rates. To increase the number of orthologs in the analysis, we calculated the average evolutionary rate for *H. sapiens* and *A. thaliana* (primary species) proteins based on *dN* values obtained from multiple pairs of evolutionary related species, while also allowing protein orthologs in some species to be missing. Specifically, we used *Gorilla gorilla*, *Pongo abelii*, *Macaa mulatta*, *Saimiri boliviensis boliviensis*, and *Mus musculus* as the secondary species for *Homo sapiens*; and *Arabidopsis halleri*, *Brassica oleracea* (cabbage), *Glycine max* (soybean), *Solanum lycopersicum* (tomato), and *Helianthus annuus* (sunflower) as the secondary species for *Arabidopsis thaliana*.

First, we estimated the evolutionary length of the branches between the primary species, *H. sapiens* and *A. thaliana*, and their corresponding secondary species. To do this, we constructed a set of proteins from the primary species that have orthologs in all secondary species. Based on these sets, we calculated the average evolutionary distance between the primary and secondary species, *k*, as 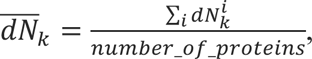 where 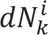 is the non-synonymous evolutionary rate of the protein *i* calculated relative to the secondary species *k*, and the sum is over all proteins, *i*, in the set. Next, for each secondary species *k*, we calculated the relative evolutionary branch length 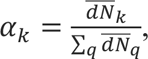 where the sum is over all secondary species, *q*. Finally, for each protein *i* with orthologs in at least one secondary species, we estimated the mean protein-specific evolutionary rate as 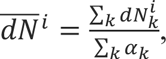 where the sum in the numerator and denominator is over the secondary species, *k*, that have orthologs of the protein *i*. The resulting protein-specific rates for orthologs in the primary species represent the fraction of non-synonymous substitutions along all evolutionary branches to the secondary species, normalized by the relative evolutionary length of the branches for which orthologs were detected.

The usage of the mean protein-specific rates, 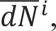 for *H. sapiens* and *A. thaliana* towards multiple secondary species increased by ∼15-20% the number of proteins with estimated evolutionary rates. Specifically, 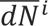 was inferred for 18634 human and 23076 *A. thaliana* proteins, compared to 16846 and 19042 proteins, respectively, when using only *Mus musculus* and *Brassica oleracea* as orthologous species (Zhang and Yang 2015).

Estimated evolutionary rates for each species are available in **Supplementary Table 5**.

The polymorphism rate for human proteins was calculated as the number of non-synonymous protein-coding SNPs with frequencies greater than 1% in the 1000 Genomes Project (Auton, et al. 2015) normalized by the protein length. The human polymorphism data was obtained from the dbSNP database (Sherry, et al. 2001) (https://www.ncbi.nlm.nih.gov/snp/).

### Expression – evolutionary Rate correlation (ER)

In the manuscript, we calculated the Expression – evolutionary Rate correlation (ER) as the Spearman’s correlation coefficient between mRNA expression levels and protein evolutionary rates. The Spearman’s correlation was used to avoid making any assumptions about the shape of the dependency between evolutionary rates and expression; the Spearman’s correlation also makes ER invariant with respect to the normalization and log-transformation of expression and evolutionary rate data. We note that all ER correlations calculated in this work are negative, indicating an anticorrelation between expression and evolutionary rate, but for visualization purposes we primarily use the correlation strength, quantified as the absolute value of the ER correlation, |ER|. Cell-type and tissue-specific ER correlations for each considered dataset are available in the **Supplementary Table 3**.

### Linear regression analysis of multi-tissue contribution to ER

The relationship between expression and evolutionary rate is non-linear (**Fig. 2**, **Supplementary fig. S2**). Therefore, to explore the joint influence of expression profiles across multiple tissues on evolutionary rates, without making any assumptions about the ER shape, we used rank-transformed expression values and rank-transformed evolutionary rates in the linear regression analysis. We then fitted the multivariable linear regression model based on expression in all tissues and compared its predictive power with the regression model based on expression in neural tissues only. The explanatory power or these regression models for different species is available in the **Supplementary Table 1**.

### Expression breadth across tissues

The breadth of expression for a gene was defined as the number of tissues in which it was expressed above a certain threshold (Park and Choi 2010). We explored several possible thresholds, namely 0, 0.1, 0.3, 1, 3, 10, 30, 100, 300, 1000 TPM. We then selected, for each animal species in the analysis, the threshold that provided the strongest Spearman’s correlation between evolutionary rate and expression breadth; the selected threshold was 10 TPM for *H. sapiens*, *M. musculus*, and *D. melanogaster*, and 30 TPM for *C. elegans*. The Spearman’s correlations between evolutionary rate and expression breadth are shown in the **Supplementary Table 1**.

### Influence of tissue-specific genes on the ER correlation

We defined the specificity of a particular gene to a tissue *k* as the z-score of its expression in tissue *k* relative to other tissues, 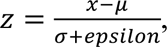 where *x* is the expression level of the gene in tissue *k*, *μ* and *σ* are the mean and standard deviation of the expression level of the gene in other tissues, and *epsilon* is a parameter equal to the minimum non-zero expression level in the transcriptome. Expression values were log10(x+1) transformed before the analysis. To investigate how well the expression of genes that are specific to a given tissue correlates with evolutionary rates, we selected various fractions of genes (ranging from 10% to 100%) with the highest specificity score to a given tissue and then used these genes to calculate the ER correlation for all tissues. This analysis was repeated for genes specific to each tissue, and the results are shown in the **Supplementary fig. S4**.

We used a similar approach to calculate the overall gene specificity to neural tissues in animals and to growing tissues in plants. In this case, the gene specificity score was defined as 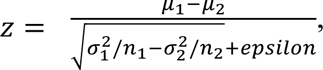 where *μ*_1_ and 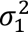 are the mean and variance of the gene expression in neural tissues of animals or in fast growing plant tissues, *μ*_2_ and 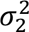 are the mean and variance of the gene expression in other tissues, *n*_1_ and *n*_2_ are the number of neural and non-neural tissues in animals or the number of fast- and slow-growing tissues in plants, and *epsilon* is a parameter with a value equal to the minimum non-zero gene expression level in the transcriptome. Expression values were log10(x+1) transformed before the analysis. Since in plants the distinction between fast- and slow-growing tissues is not binary, we classified 20% of the tissues with the highest expression of growth markers as fast-growing and 20% of the tissues with the lowest expression of growth markers as slow-growing. To evaluate the influence of neural-specific genes in animals and fast-growth-specific genes in plants on the ER correlation, we removed 10% of the most neural- or fast-growth genes from the analysis and recalculated the ER correlations. The results of this analysis are shown in the **Supplementary Table 3**.

### Influence of the number of expressed genes on the ER correlations

The number of expressed genes varies across tissues. To explore how this variability might contribute to the differences in the strength of tissue-specific ER we equalized the number of expressed genes across tissues. To that end, for each multi-tissue transcriptome, we identified the tissue with the minimum number of non-zero expressed genes, *n*. For each other tissue, we then sorted the genes based on their expression levels and selected *n* genes, starting with the most highly expressed, and set expression values for the other gens to zero. We then recalculated the ER correlations across tissues using these modified expression profiles and compared them to the original ER correlations. The results of this analysis are shown in the **Supplementary Table 3**.

### Expression of essential genes across tissues

We used the data describing the fitness effects of ∼7000 knockouts for mouse genes obtained from the International Mouse Phenotyping Consortium (IMPC) database available for download on April 18^th^ 2022 (Koscielny, et al. 2014) [https://www.mousephenotype.org/about-impc/]. We defined genes to be essential if their knockout viability phenotypes were classified in the database as “lethal”. We calculated the fraction of essential genes expressed (at the level ≥1 TPM) in the mouse brain at different developmental stages; the results were not sensitive to the selected expression threshold.

### Gene set enrichment analysis (GSEA)

Gene ontology (GO) annotations were obtained from the Molecular Signatures Database MSigDB:C5 v7.4 (release date March 2021) (Liberzon, et al. 2011) for mouse genes, from the Arabidopsis Information Resource (TAIR) database (release date April 1^st^ 2021) (Berardini, et al. 2015) for Arabidopsis genes, and from the AmiGo browser (release date February 1^st^ 2021) (Carbon, et al. 2009) for fly, corn and soybean genes.

For GSEA analyses of single-cell transcriptomes, we maintained the consistency in the resolution of expression profiles across cell-type clusters by retaining the same number of non-zero expressed genes in each cell-type cluster. To achieve this, we sorted genes by their expression for each cell-type and retained the top 10,000 most highly expressed genes for mouse and 7,000 genes for *D. melanogaster*. Expression of other genes was set to zero. We note that GSEA analyses performed on uncorrected expression data yielded very similar sets of upregulated GO terms (**Supplementary Table 2**).

We used the gene set enrichment analysis to identify the functional roles of genes with upregulated expression in cell-types exhibiting stronger ER correlations. To this end, we first calculated for each gene the Pearson’s correlation coefficient between its expression across different cell-type clusters and the strength of the cell-type-specific ER, *r*(*Expression*, |*ER*|). We then ranked the genes based on the strength of this correlation and performed a pre-ranked GSEA analysis (Subramanian, et al. 2005), using the “prerank” module of GSEAPY (Fang 2020), to identify GO terms enriched among genes that associated with stronger ER. Following the default settings, we used in the GSEA analysis GO terms containing ≥ 15, but ≤ 2000 genes.

To verify the robustness of our results, we also utilized an alternative procedure to identify the association between gene expression and the ER strength. Specifically, we sorted cell-types by the strength of ER and calculated for each gene the difference in its average expression levels between the top 10% and bottom 90% (or top 50% and bottom 50%) of the sorted cell-types. We then performed a pre-ranked GSEA analysis based on the calculated differential expression values.

We further investigated the GSEA results in mouse to understand whether the association between the ER strength and expression of genes from synaptic and other enriched GO terms was due to the direct influence of these genes on the ER correlation. To address this question, we removed all genes annotated with (i) the Synapse GO term (GO:0045202) or (ii) with any of the significantly enriched GO terms (at FDR < 0.05). For each cell-type, we then recalculated the Spearman’s correlation between evolutionary rates and expression using the remaining genes, *ER*_*noSynapse*_ and *ER*_*noEnric*ℎ*ed*_, respectively. Next, we repeated the pre-ranked GSEA analysis, but this time the gene ranking (for all genes, including synaptic and associated with other enriched GO terms) was based on the Pearson’s correlation of gene expression with *r*L*Expression*, M*ER*_*noSynapse*_MN or *r*(*Expression*, |*ER*_*noEnric*ℎ*ed*_|).

The results of the GSEA analyses for all animals’ and plants’ datasets and corresponding controls are available in the **Supplementary Table 2**.

### Hierarchical clustering of expression samples

We performed the hierarchical clustering of expression samples for each plant species using the ‘hclust’ function from the R package ‘stats’ (R Core Team 2022). The distance between samples was calculated as one minus the squared Pearson’s correlation coefficient between corresponding expression profiles. We used the ‘ward.D2’ agglomeration method for corn and the ‘complete’ method for Arabidopsis and soybean.

### Markers for growth stages in plants

Marker genes for the division, elongation, and differentiation growth stages in Arabidopsis were taken from Huang et al. (Huang and Schiefelbein 2015). For corn and soybean, we used orthologs of the Arabidopsis marker genes as the markers for the corresponding growth stages. Ortholog annotations were obtained from the Ensemble database (Cunningham, et al. 2022) via BioMart (Kinsella, et al. 2011), using the ‘one2any’ homology and confidence level ‘1’. The average expression of genes from each growth stage was calculated after log10(x+1) transformation of expression values. The final lists of marker genes for each plant species are available in **Supplementary Table 6**.

### Collection of data on enzymes’ catalytic rates

We extracted all available data on *k*_*cat*_ and *k*_*cat*_/*K*_*m*_ from the Brenda (Chang, et al. 2021) (version of September 3^rd^ 2019) and Sabio-RK (Wittig, et al. 2018) (version of January 21^st^ 2019) databases. Specifically, we downloaded from Brenda (https://www.brenda-enzymes.org/download.php) an easy-to-parse text file containing catalytic rates along with additional information about the entries, such as reaction EC numbers, protein Uniprot IDs, protein names and types, source organisms, and measurement temperatures. For Sabio-RK, we used the provided URL request interface to automatically download all kinetic constants and entries’ information. In addition, we obtained data on *k*_*cat*_for *E. coli* enzymes from the previously published and manually curated dataset (Davidi, et al. 2016). We then applied several filters to exclude mutant enzymes, enzymes with macromolecular substrates or involved in transmembrane transport, and multi-functional enzymes that catalyze multiple reactions with EC numbers different by more than the last digit. In cases where multiple values of *k*_*cat*_ or *k*_*cat*_/*K*_*m*_ were available for a given enzyme, for example due to measurement with different homologous substrates or measurements available in different publications, we used the highest values of experimentally obtained kinetic constants.

For the final sets on enzymes from *H. sapiens*, *A. thaliana*, and *E. coli* we manually curated the references where *k*_*cat*_, *k*_*cat*_/*K*_*m*_, and corresponding *k*_*cat*_^*max*^, and 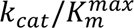 (described in the next section) were published. We observed some discrepancies between the values given in the original publications and in the corresponding entries in the Brenda or Sabio-RK databases. The most common errors were due to incorrect transfer of units, for example the usage of *mM* instead of *μM* or *s*^-1^ instead of *min*^-1^. All such cases were corrected in our dataset. Information on all manual corrections is available in the **Supplementary Table 7**.

The complete dataset of catalytic rates for the enzymes used in this work is available in the **Supplementary Table 7**. Data on catalytic rates for enzymes from *H. sapiens*, *A. thaliana*, and *E. coli*, combined with their evolutionary rates and expression values are available in the **Supplementary Table 8**.

### Normalization of catalytic rates to estimate functional optimization

Following the previous approach (Davidi, et al. 2018), to estimate the level of functional optimization of the considered enzymes we calculated the normalized catalytic rates, *k*_*cat*_^*norm*^ or 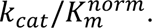 To that end, we divided the kinetic constants, *k*_*cat*_ and *k*_*cat*_/*K*_*m*_, by the highest constants known for the corresponding reaction classes, *k*_*cat*_^*max*^ and 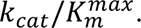 This normalization procedure allowed us to account for a substantial heterogeneity of kinetic constant values across different reactions classes due to the diverse chemistries of the catalyzed reactions. To perform the normalization, we identified for each reaction class, i.e., enzymes sharing all four digits of the EC classification, the highest catalytic rates measured across all database entries (including mutants and multifunctional enzymes), *k*_*cat*_^*max*^ or 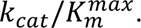 To ensure accurate estimates of the maximum catalytic rate for a given reaction class, we included in the analysis only enzymes for which experimental measurements of catalytic rates were available for at least 15 different unique enzymes in the same reaction class (EC number); the results were not very sensitive to the value of this parameter.

### Temperature correction of the measured catalytic rates

The majority of the catalytic rate constants in our dataset were measured at temperatures in the range of 20°C to 40°C, with the median ∼30°C. However, approximately 6% of the available kinetic measurements have been performed at higher temperatures, up to 100°C. Since the rates of catalyzed biochemical reactions are temperature-dependent according to the Arrhenius’ law, we applied a temperature correction to estimate the catalytic rates of the corresponding enzymes at 30°C. We note that the exact dependence of the reaction rate on temperature depends on the activation energy, which is specific to each catalyzed chemical reaction. However, a previous systematic analysis of the temperature-dependent acceleration of biochemical reactions, catalyzed by more than 150 enzymes, demonstrated (Elias, et al. 2014) that on average the values of *k*_*cat*_and *k*_*cat*_/*K*_*m*_ increased by a factor of 1.8 for each 10°C increase in temperature. To account for this trend, we scaled down all *k*_*cat*_ and *k*_*cat*_/*K*_*m*_ values experimentally measured at temperatures *T* > 40°C using the following equation: *k*(30°C) = *k*(*T*) · 1.8^(*T*–30)⁄10^, where *k* is the catalytic constant and *T* is the measurement temperature; we did not apply any adjustments to the catalytic rate constants measured at *T* < 40°C. In cases where the measurement temperature was not explicitly stated in the publication, we assumed that the measurements were performed at ambient temperature and did not apply any temperature correction.

We note that temperature correction only affected the functional efficiency of 21 enzymes (9%) in the final sets for Human, Arabidopsis, and E. coli. Moreover, functional efficiencies based on the uncorrected data showed very similar correlations with protein evolutionary rates and expression to the temperature-corrected data (**Supplementary Table 9**).

### Multi-member EC classes

Species’ genomes often encode several distinct enzymes that catalyze the same EC reaction. Among the sets of enzymes with estimated functional efficiency most reaction classes were represented only by one or two different proteins. However, for *H. sapiens* and *A. thaliana*, several EC classes were represented by multiple enzymes. These enzymes usually had different expression levels, evolutionary rates, and catalytic efficiencies (**Supplementary Table 8**). However, to assess the potential influence of multi-member EC classes on the KR correlation, we randomly subsampled the enzyme from the multi-member EC classes and retained in each sampled trial only two different enzymes from each EC class with more than two protein members. For each of the 10,000 random trials, we recalculated the KR correlation and the fraction of the ER correlation mediated by the KR correlation. Although this procedure reduced the number of enzymes by about 1/3, the KR correlation was significant in >99% of the reduced samples, and the median of the KR correlation coefficients and the fractions of the ER explained by functional efficiency were similar to the results obtained from the complete enzyme sets (**Supplementary Table 9**).

### Unique and shared contributions of protein optimality and expression to explaining protein evolutionary rates

We used semi-partial correlations (Abdi 2007) to investigate the unique and shared contributions of the two independent variables, i.e., protein functional optimality and expression, to explaining the variance of the dependent variable, protein evolutionary rates. To that end, the unique effect of one independent variable *x* on the dependent variable *z* was calculated as the squared coefficient of the semi-partial correlation between them while controlling for the other independent variable, *y*, 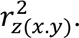 Likewise, the unique effect of the independent variable y on the dependent variable *z* was calculated as 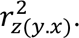 The shared effect was calculated as the difference between the squared coefficient of the ordinary bivariate correlation, 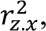 and the squared coefficient of the semi-partial correlation 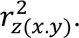 We used the function “partial_cor” with the method “spearman” from the Python package “pingouin” to calculate semi-partial correlations (Vallat 2018).

