## Supplemental_Figures for "Functional optimization in distinct tissues and conditions constrains the rate of protein evolution"

### SUPPLEMENTARY FIGURES

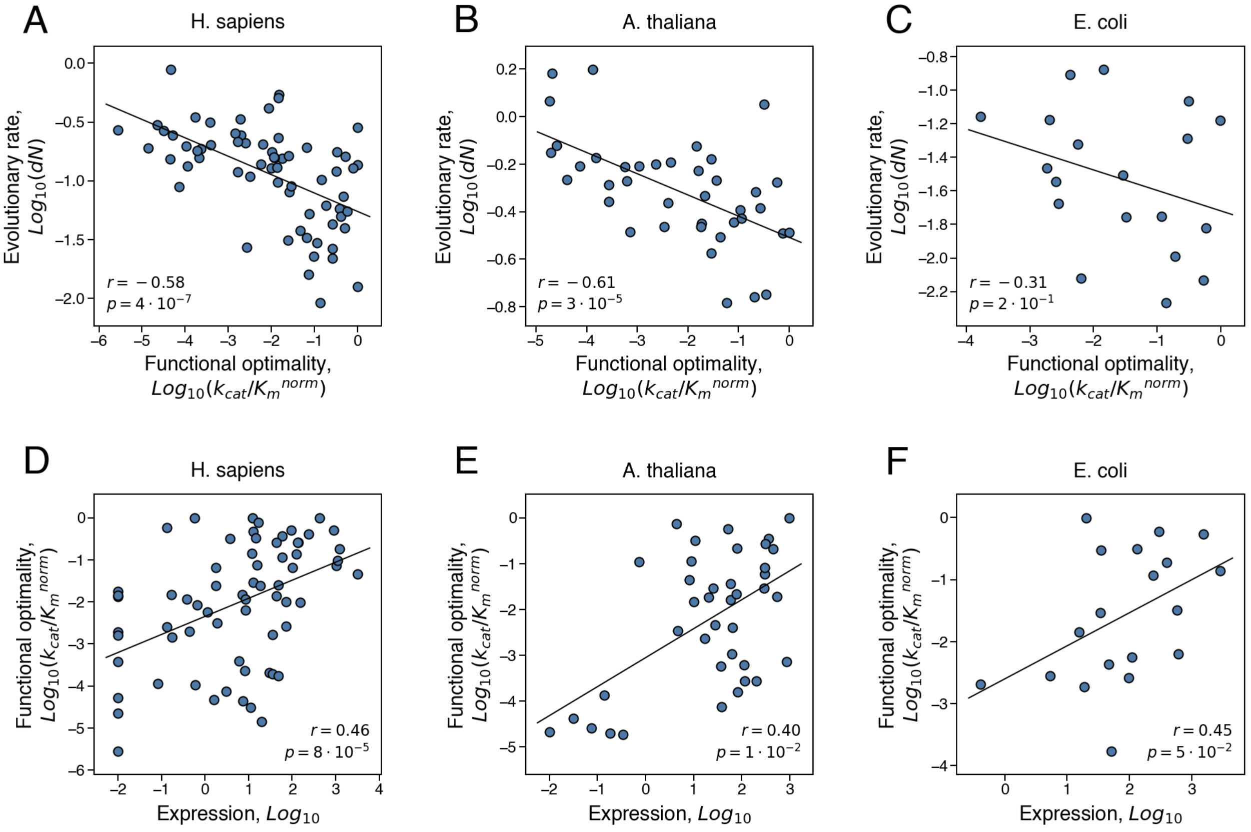


**Supplementary fig. S1. The correlations between protein functional optimality, quantified as $k_{cat}/K_M^{norm}$, expression, and the rate of protein evolution.** (**A, B, C**) The correlation between protein functional optimality and the rate of protein evolution (the KR correlation), and (**D, E, F**) the correlation between expression level and protein functional optimality (the EK correlation). Each point in the figures represents an enzyme from (**A, D**) *H. sapiens* (n=66), (**B, E**) *A. thaliana* (n=39), and (**C, F**) *E. coli* (n=19). Protein functional optimality was estimated using the normalized second-order kinetic constant, $k_{cat}/K_M^{norm}$, which quantifies how quickly an enzyme performs the reaction in low substrate conditions relative to the maximal second-order kinetic constant measured for the same reaction class. Evolutionary rate, *dN*, was calculated as the number of non-synonymous substitutions accumulated during the divergence of closely related orthologs per non-synonymous site. For multicellular species, the expression in the tissue with the strongest ER for enzymes was used (the brain basal ganglia for *H. sapiens* and the seedling root for *A. thaliana*). Spearman's correlation coefficients and p-values are shown in each figure.

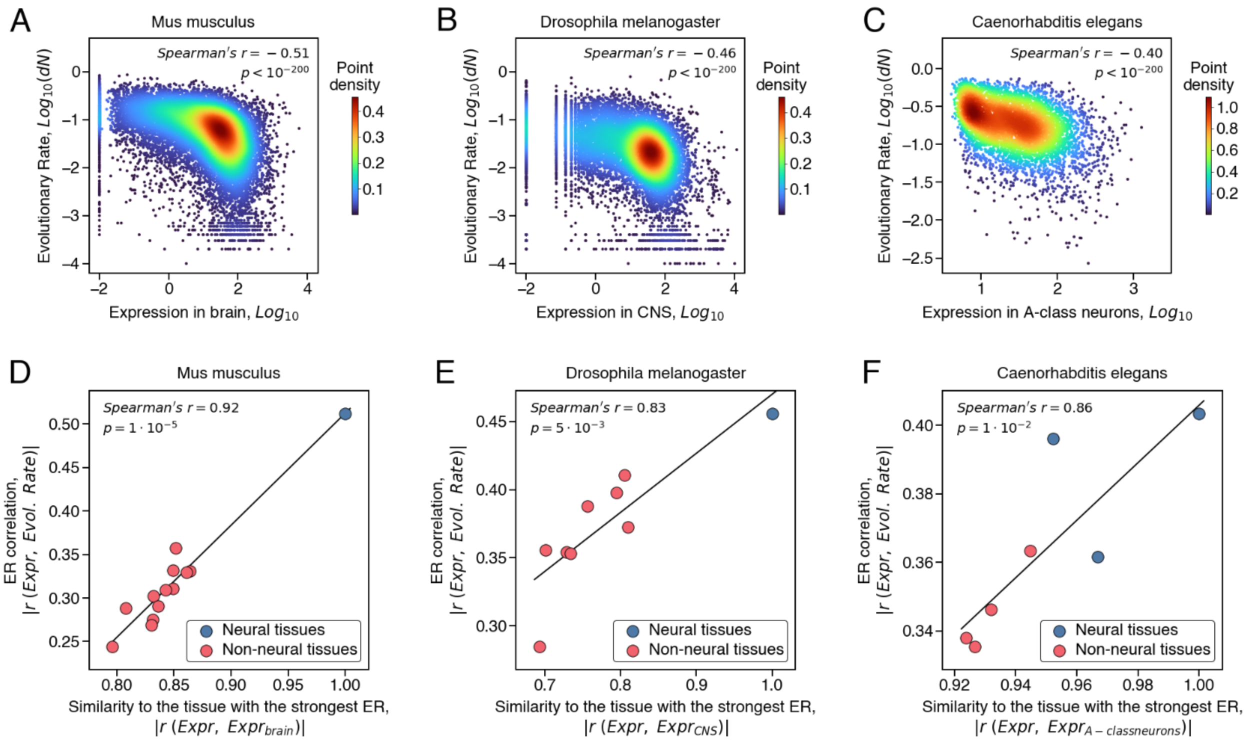


**Supplementary fig. S2. The relationship between tissue-specific expression and the rate of protein evolution in different animals.** (**A**, **B**, **C**), The correlation between gene expression in the neural tissues and evolutionary rates of the corresponding proteins in (**A**) *M. musculus* (n=17249), (**B**) *D. melanogaster* (n=12545), and (**C**) *C. elegans* (n=5384). Evolutionary rates, *dN*, were calculated as the number of non-synonymous substitutions accumulated during the divergence of closely related orthologs per non-synonymous site (see Methods). Each point on the plot represents a protein and the colors represent the point density. (**D**, **E**, **F**), The correlation between ER values across tissues of (**D**) *M. musculus*, (**E**) *D. melanogaster*, and (**F**) *C. elegans* and the similarity of tissues' genes expression to the neural tissue with the strongest ER, which is the brain, central nervous system, and A-class neuron, respectively. The similarity between tissues' expression profiles was quantified using the Spearman's correlation. Blue points represent neural tissues, and red points represent non-neural tissues. The linear regression and Spearman's correlation coefficients were calculated based on all tissues (n=13, 9, and 7, respectively).

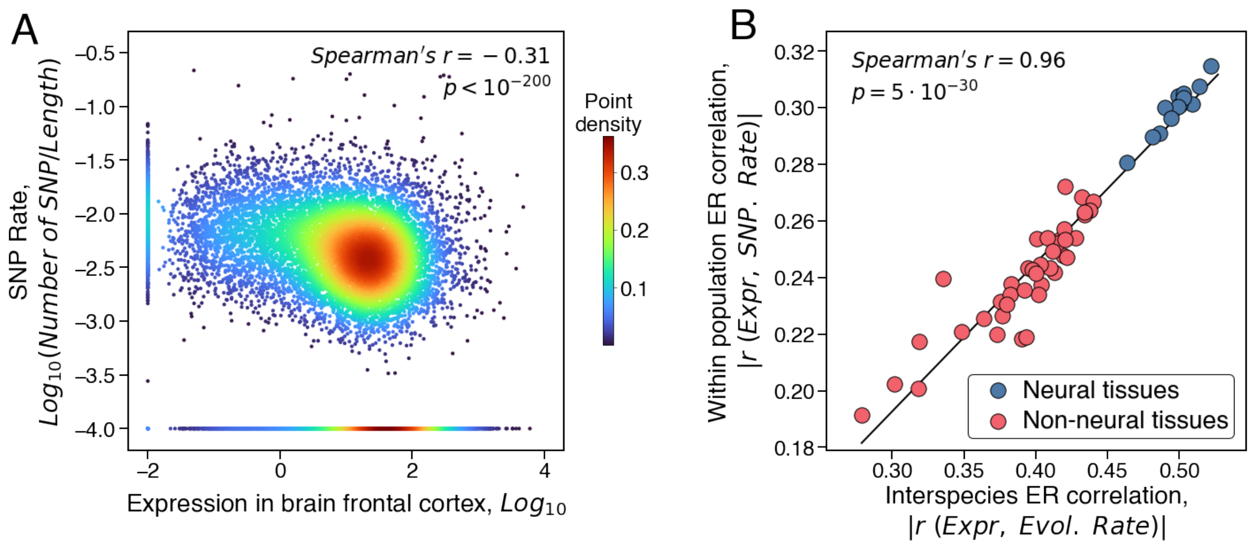


**Supplementary fig. S3. The relationship between tissue-specific expression and the frequency of polymorphisms in human genes.** (**A**) The correlation between gene expression in the human brain frontal cortex and the frequency of single nucleotide polymorphisms (SNP) in human genes; the polymorphism frequency was calculated as the number of SNPs (with population frequencies greater than 1%) per protein, normalized by the protein length. Each point in the plot represents a human protein (n=15509), and the colors represent the point density. **(B)** The correlation between ER values across human tissues calculated using different measures of protein evolutionary rates: the interspecies divergence of protein sequences (X-axis) and the SNP frequencies (Y-axis). Blue points represent neural tissues (n=13), and red points represent non-neural tissues (n=40). The linear regression and the Spearman's correlation coefficient were calculated based on all 53 tissues.

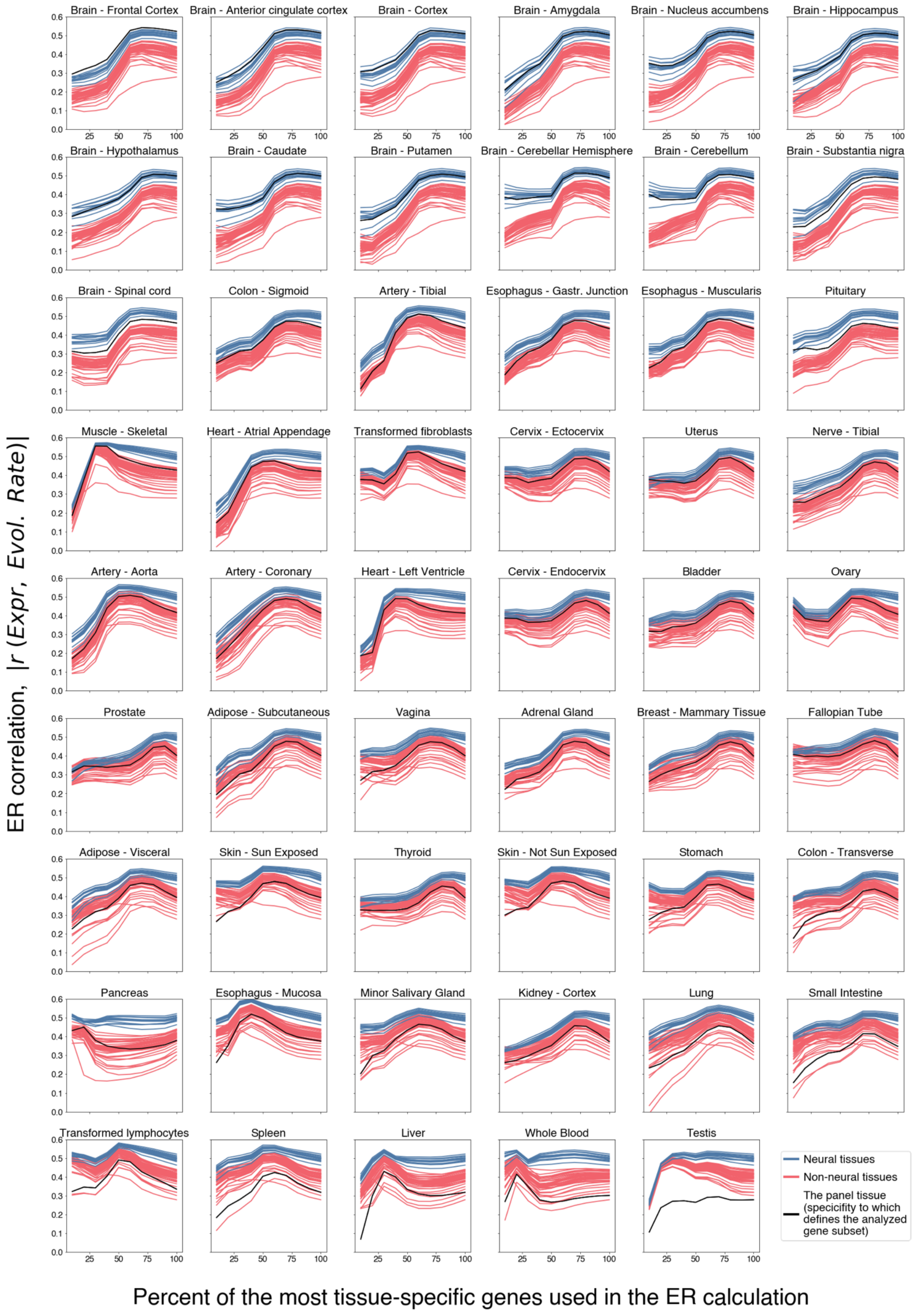

Brain - Frontal Cortex
Brain - Anterior cingulate cortex
Brain - Cortex
Brain - Amygdala
Brain - Nucleus accumbens
Brain - Hippocampus
Brain - Hypothalamus
Brain - Caudate
Brain - Putamen
Brain - Cerebellar Hemisphere
Brain - Cerebellum
Brain - Substantia nigra
Brain - Spinal cord
Colon - Sigmoid
Artery - Tibial
Esophagus - Gastr. Junction
Esophagus - Muscularis
Pituitary
Muscle - Skeletal
Heart - Atrial Appendage
Transformed fibroblasts
Cervix - Ectocervix
Uterus
Nerve - Tibial
Artery - Aorta
Artery - Coronary
Heart - Left Ventricle
Cervix - Endocervix
Bladder
Ovary
Prostate
Adipose - Subcutaneous
Vagina
Adrenal Gland
Breast - Mammary Tissue
Fallopian Tube
Adipose - Visceral
Skin - Sun Exposed
Thyroid
Skin - Not Sun Exposed
Stomach
Colon - Transverse
Pancreas
Esophagus - Mucosa
Minor Salivary Gland
Kidney - Cortex
Lung
Small Intestine
Transformed lymphocytes
Spleen
Liver
Whole Blood
Testis
Neural tissues
Non-neural tissues
The panel tissue (specificity to which defines the analyzed gene subset)
ER correlation, |r (Expr, Evol. Rate)|
Percent of the most tissue-specific genes used in the ER calculation

**Supplementary fig. S4. Expression-evolutionary Rate (ER) correlations calculated for subsets of human genes with various expression specificities to different tissues.** Each panel in the figure describes the effect of using only genes specific to a particular tissue (labeled above each panel) on the strength of the ER correlation calculated based on gene expression in all human tissues. Each line in a panel represents ER calculated for a human tissue using a certain fraction of genes specific to the panel tissue. The Y-axis shows the strength of the ER correlation, and the X-axis shows the fraction of the most specific genes from the panel tissue used to calculate ER. Specificity of a gene to a given tissue was defined as the z-score characterizing the difference between the gene's expression level in that tissue and the mean expression level across all tissues, normalized by the standard deviation of the gene expression levels across all tissues. The blue lines in the plots represent ER calculated based on expression in human neural tissues, the red lines represent ER calculated based on expression in non-neural tissues, and the black line represents ER calculated based on expression in the panel tissue.

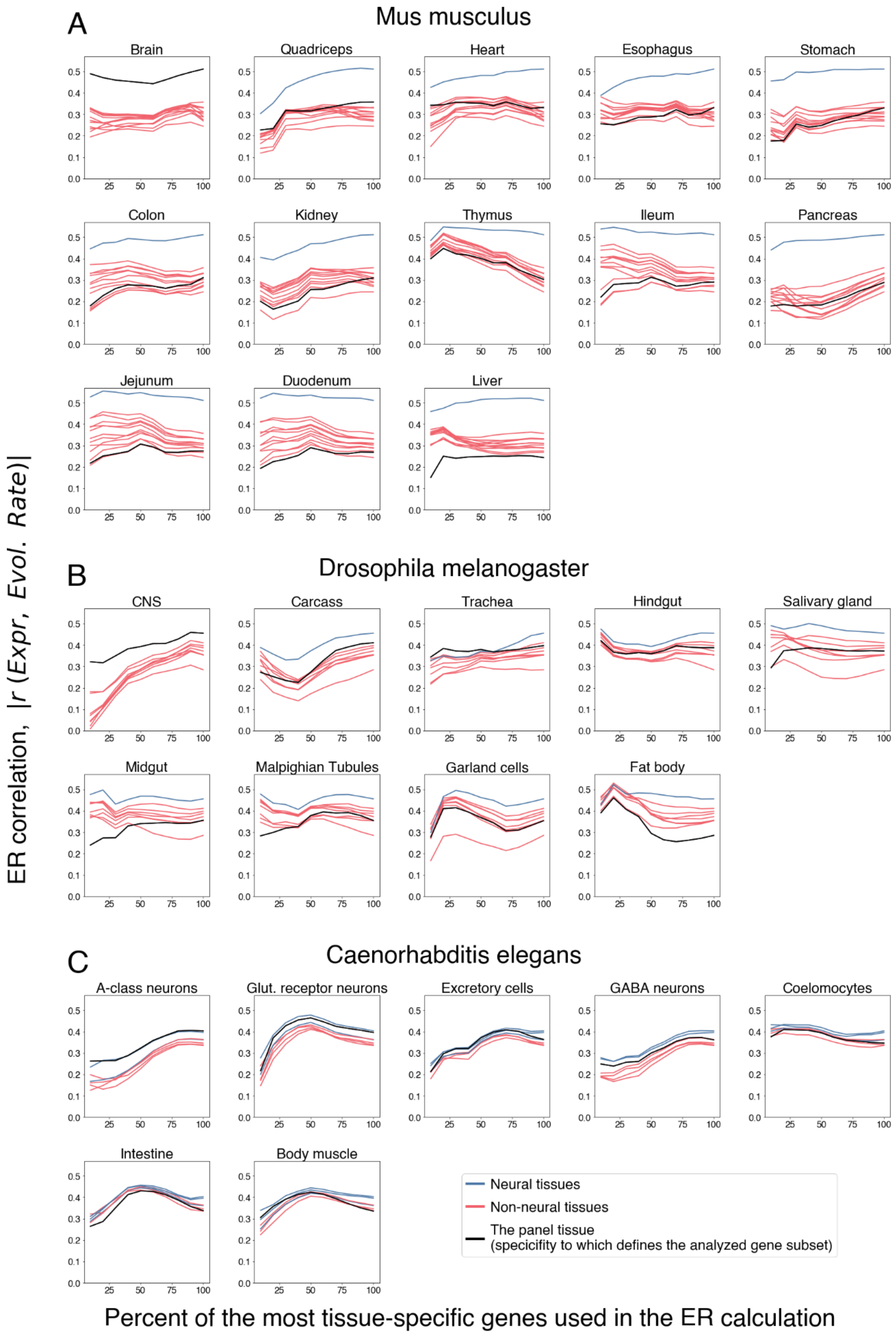

A
Mus musculus
Brain
Quadriceps
Heart
Esophagus
Stomach
Colon
Kidney
Thymus
Ileum
Pancreas
Jejunum
Duodenum
Liver
B
Drosophila melanogaster
CNS
Carcass
Trachea
Hindgut
Salivary gland
Midgut
Malpighian Tubules
Garland cells
Fat body
C
Caenorhabditis elegans
A-class neurons
Glut. receptor neurons
Excretory cells
GABA neurons
Coelomocytes
Intestine
Body muscle
0.0
0.1
0.2
0.3
0.4
0.5
25
50
75
100
Neural tissues
Non-neural tissues
The panel tissue
(specificity to which defines the analyzed gene subset)
ER correlation, |r (Expr, Evol. Rate)|
Percent of the most tissue-specific genes used in the ER calculation

**Supplementary fig. S5. Expression-evolutionary Rate (ER) correlations in different animals calculated for subsets of genes with various expression specificities to different tissues.** The analysis was conducted for (**A**) *M. musculus*, (**B**) *D. melanogaster*, and (**C**) *C. elegans*. Each panel in the figure describes the effect of using only genes specific to a particular tissue (labeled above each panel) on the strength of the ER correlation calculated based on gene expression in all tissues. Each line in a panel represents ER calculated for a tissue using a certain fraction of genes specific to the panel tissue. The Y-axis shows the strength of the ER correlation, and the X-axis shows the fraction of the most specific genes from the panel tissue used to calculate ER. Specificity of a gene to a given tissue was defined as the z-score characterizing the difference between the gene's expression level in that tissue and the mean expression level across all tissues, normalized by the standard deviation of the gene expression levels across all tissues. The blue lines in the plots represent ER calculated based on expression in neural tissues, the red lines represent ER calculated based on expression in non-neural tissues, and the black line represents ER calculated based on expression in the panel tissue.

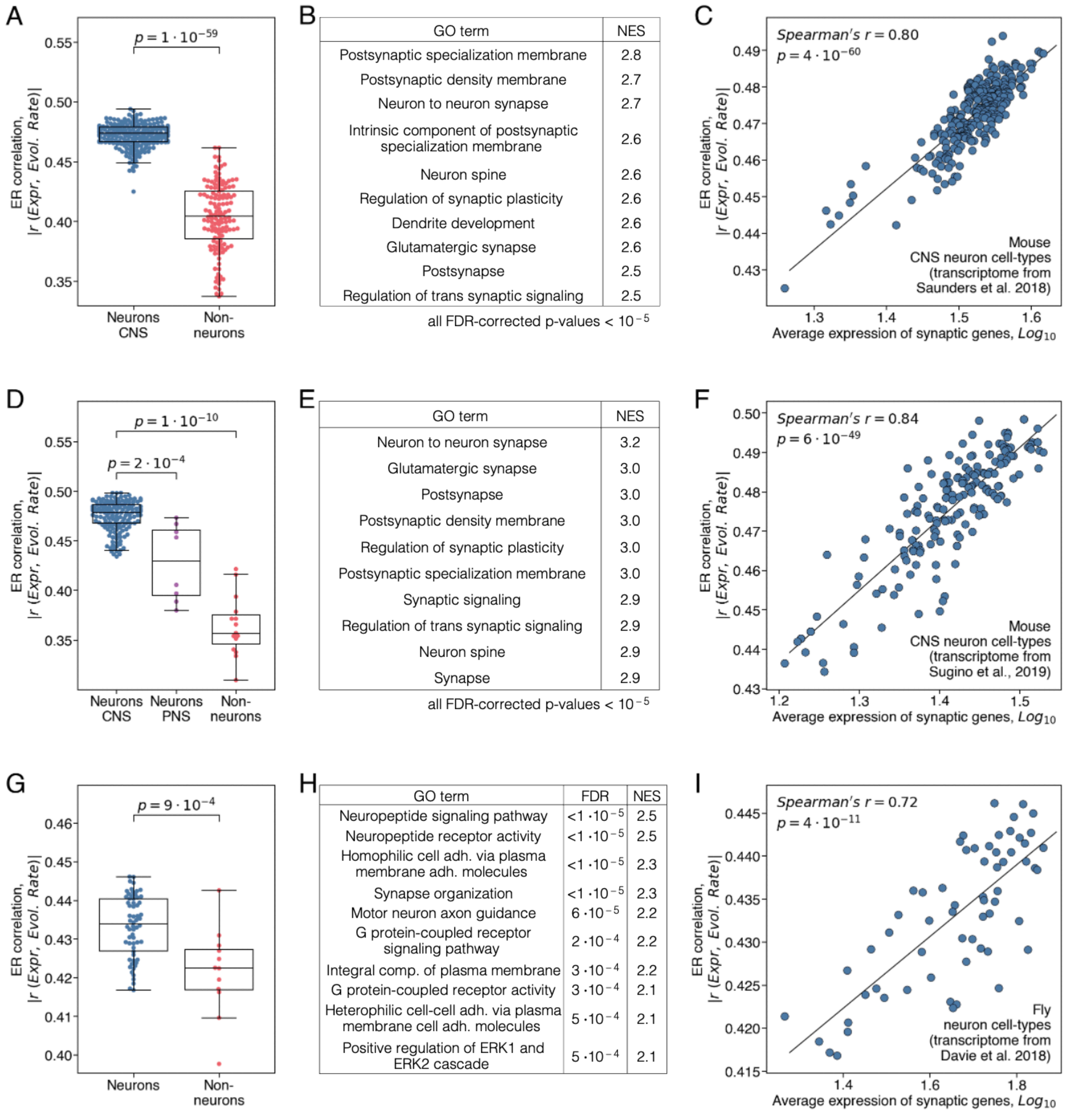


**Supplementary fig. S6. Functional properties of animals' brain cells that are associated with stronger ER correlations.** (**A**, **D**, **G**) The strength of ER correlations across different cell types in the mouse nervous system (**A**) based on the single-cell sequencing (Saunders, et al. 2018) (n=404) and (**B**) bulk mRNA sequencing (Sugino, et al. 2019) (n=202) transcriptomes, and (**C**) in the brain of *D. melanogaster* (Sugino, et al. 2019) (n=74). CNS neurons are shown in blue, PNS neurons in purple, and non-neuronal brain cells in red. The box plots show the median, the upper and lower quartiles of the ER strength, and the whiskers show the minimum and maximum values excluding outliers; P-values were calculated using the Mann-Whitney U-test. (**B**, **E**, **H**) The Gene Ontology (GO) terms associated with stronger ER in mouse CNS neurons obtained by (**B**) single-cell or (**E**) bulk sequencing and (**H**) in *D. melanogaster* neurons. FDR-corrected P-values and normalized enrichment scores (NES) are shown for the 10 top GO terms ranked by their NES. The complete list of significantly associated GO terms is provided in the **Supplementary Table 2**. (**C**, **F**, **I**) The correlation between the average expression of genes from the Synapse GO term (GO:0045202) for mouse or from the Synapse organization GO term (GO:0050808) for fly and the cell-type specific ER strength. Each point represents either (**C**) a mouse CNS neuron type from single-cell sequencing dataset (n=268), or (**F**) a mouse CNS neuron type from bulk sequencing dataset (n=179) dataset, or (**I**) a *D. melanogaster* brain neuron type (n=62).

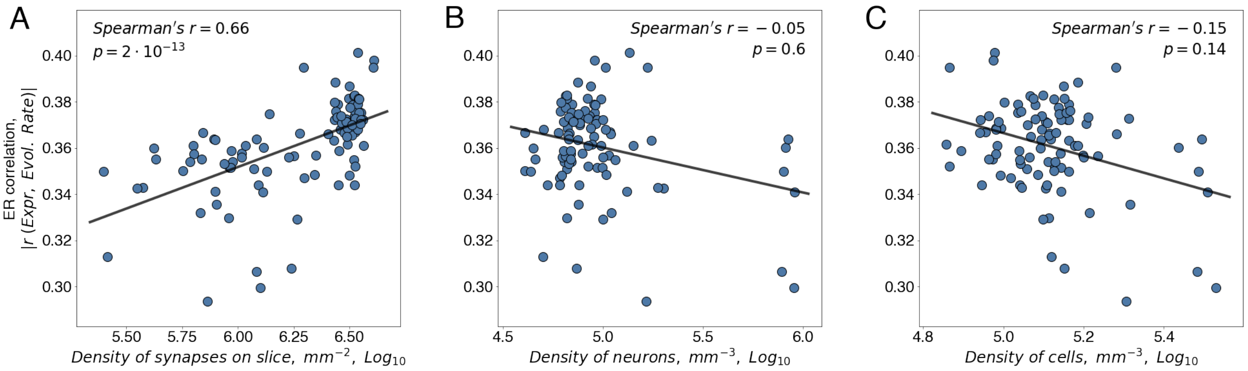


**Supplementary fig. S7. The correlation between the strength of ER and synaptic and cell densities across the mouse brain regions.** The relationship between and the strength of ER and (**A**) the synaptic density (Zhu, et al. 2018), (**B**) the density of neurons (Erö, et al. 2018), and (**C**) the density of all cells (Murakami, et al. 2018) in each region of the mouse brain. Figure points represent 96 mouse brain regions for which expression and the cellular and synaptic densities were previously measured.

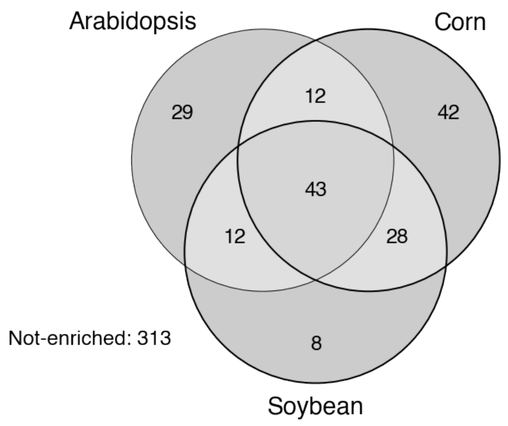


**Supplementary fig. S8. The overlap between GO functional terms that are associated with stronger ER correlations in the considered plant species.** The Venn diagram shows the overlap of GO terms associated (FDR<0.05) with stronger ER in the gene set enrichment analysis of the considered plant species (corn, Arabidopsis, and soybean). The two-way overlaps represent the number of GO terms associated with stronger ER in the pairs of corresponding species. The three-way overlap represents the number of GO terms associated with stronger ER in all three plants. The probability of observing 43 GO terms in the 3-way overlap by chance, considering that 121, 92, and 95 GO terms are associated with the ER strength in individual plant species, $p = 10^{-37}$, was calculated using the R package "SuperExactTest" (Wang, et al. 2015).

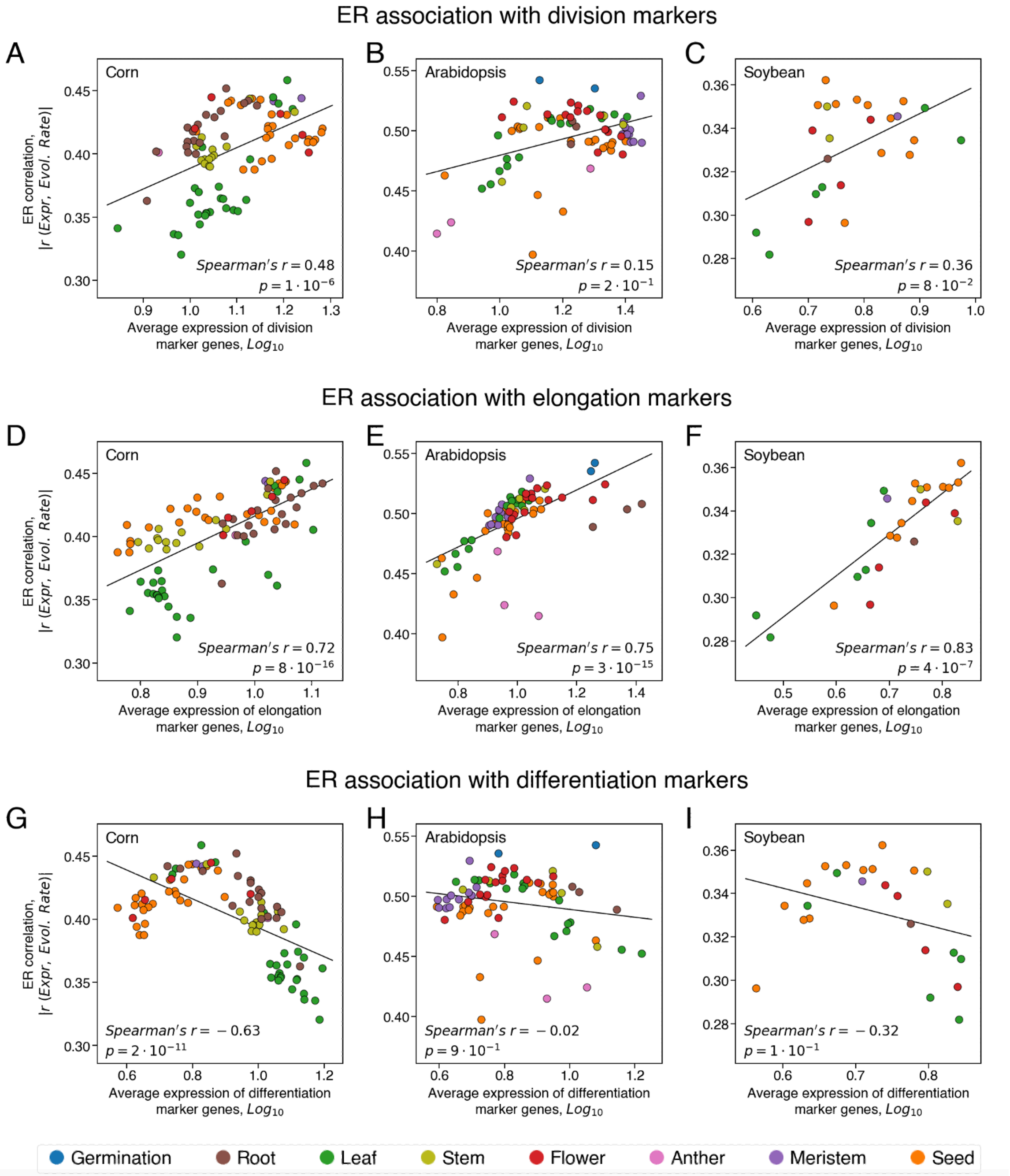


**Supplementary fig. S9. The correlation between ER and the expression level of growth-related gene markers in the considered plant species.** The correlation between ER and the average expression of cell division markers (**A-C**), elongation markers (**D-F**), and differentiation markers (**G-I**) in plant tissues. Each point in the figure represents a tissue from (**A**, **D**, **G**) corn (n=92), (**B**, **E**, **H**) Arabidopsis (n=79), and (**F**, **F**, **I**) soybean (n=25). The point colors represent the major plant tissue types as specified in the legend. For Arabidopsis, the gene markers of division, elongation, and differentiation previously identified in the analysis of Arabidopsis root cells from the corresponding growth stages (Huang and Schiefelbein 2015) were used. For corn and soybean, the orthologs of the corresponding Arabidopsis genes were used as gene markers.

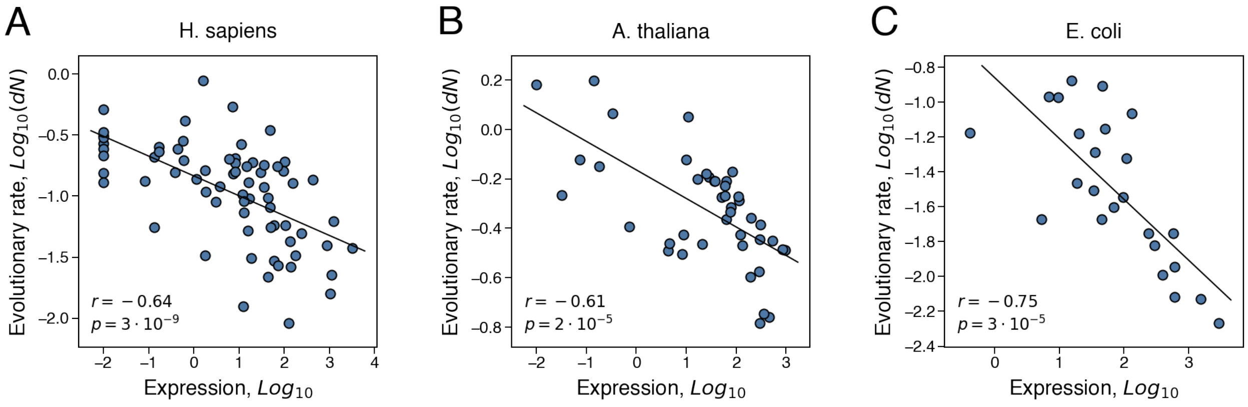


**Supplementary fig. S10. The ER correlation calculated for enzymes with estimated functional optimality ($k_{cat}^{norm}$) values.** The ER correlation between the rate of evolution for enzymes (Y-axis) and expression level of the corresponding genes (X-axis). Each point in the figure represents an enzyme from (**A**) *H. sapiens* (n=70), (**B**) *A. thaliana* (n=42), and (**C**) *E. coli* (n=24) with estimated $k_{cat}^{norm}$ value. Evolutionary rates, *dN*, were calculated as the number of non-synonymous substitutions accumulated during the divergence of closely related orthologs per non-synonymous site. For multicellular species, the expression in the tissue with the strongest ER for enzymes was used (the brain basal ganglia for *H. sapiens* and the seedling root for *A. thaliana*).

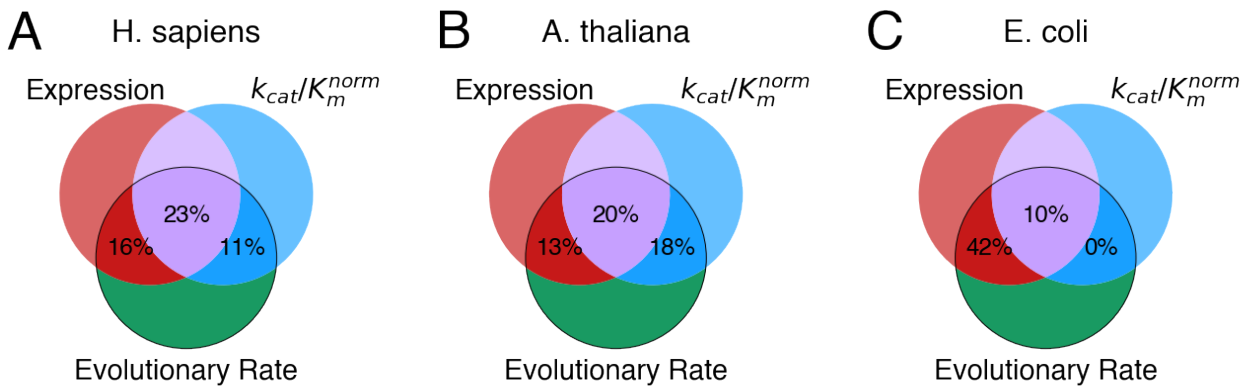


**Supplementary fig. S11. The fraction of the evolutionary rate variance explained by protein functional optimality, quantified as $k_{cat}/K_M^{norm}$, and expression.** The Venn diagram shows for (**A**) *H. sapiens*, (**B**) *A. thaliana*, and (**C**) *E. coli*, the fractions of the evolutionary rate variance explained by expression and protein functional optimality; functional optimality was quantified using the normalized kinetic constant $k_{cat}/K_M^{norm}$. The two-way intersections (red and blue) represent the unique contributions of expression and functional optimality, respectively, and three-way intersection (purple) represents the shared contribution of these two factors. For multicellular species, the expression in the tissue with the strongest ER for enzymes was used (the brain basal ganglia for *H. sapiens* and the seedling root for *A. thaliana*). The unique and shared contributions were estimated using semi-partial correlations (see Methods).
